# Mechanism of cellular production and *in vivo* seeding effects of hexameric β-amyloid assemblies

**DOI:** 10.1101/2020.12.23.424094

**Authors:** Céline Vrancx, Devkee M Vadukul, Sabrina Contino, Nuria Suelves, Ludovic D’Auria, Florian Perrin, Vincent Van Pesch, Bernard Hanseeuw, Loïc Quinton, Pascal Kienlen-Campard

## Abstract

**Background:** The β-amyloid peptide (Aβ) plays a key role in Alzheimer’s disease. After its production by catabolism of the amyloid precursor protein (APP) through the action of presenilin 1 (PS1)- or presenilin 2 (PS2)-dependent γ-secretases, monomeric Aβ can assemble in oligomers. In a pathological context, this eventually leads to the formation of fibrils, which deposit in senile plaques. Many studies suggest that Aβ toxicity is related to its soluble oligomeric intermediates. Among these, our interest focuses on hexameric Aβ, which acts as a nucleus for Aβ self-assembly.

**Methods:** Biochemical analyses were used to identify hexameric Aβ in a wide range of models; cell lines, cerebrospinal fluid from cognitively impaired patients and transgenic mice exhibiting human Aβ pathology (5xFAD). We isolated this assembly and assessed both its effect on primary neuron viability *in vitro*, and its contribution to amyloid deposition *in vivo* following intracerebral injection. In both cases, we used wild-type mice (C57BL/6) to mimic an environment where hexameric Aβ is present alone and 5xFAD mice to incubate hexameric Aβ in a context where human Aβ species are pre-existing. Using CRISPR-Cas9, we produced stable *knockdown* human cell lines for either PS1 or PS2 to elucidate their contribution to the formation of hexameric Aβ.

**Results:** In WT mice, we found that neither *in vitro* or *in vivo* exposure to hexameric Aβ was sufficient to induce cytotoxic effects or amyloid deposition. In 5xFAD mice, we observed a significant increase in neuronal death *in vitro* following exposure to 5μM hexameric Aβ, as well as a 1.47-fold aggravation of amyloid deposition *in vivo*. At the cellular level, we found hexameric Aβ in extracellular vesicles and observed a strong decrease in its excretion when PS2 was knocked down by 60%.

**Conclusions:** Our results indicate the absence of cytotoxic effects of cell-derived hexameric Aβ by itself, but its capacity to aggravate amyloid deposition by seeding other Aβ species. We propose an important role for PS2 in the formation of this particular assembly in vesicular entities, in line with previous reports linking the restricted location of PS2 in acidic compartments to the production of more aggregation-prone Aβ.

## Background

The β-amyloid peptide (Aβ) is the major constituent of the senile plaques, a typical histological hallmark of Alzheimer’s disease (AD). This peptide is produced by the amyloidogenic catabolism of the amyloid precursor protein (APP) [1]. APP undergoes a first cleavage by β-secretase, producing a C-terminal fragment (βCTF), which in turn is cleaved by γ-secretase to generate the intracellular domain of APP (AICD) and Aβ. To note, different isoforms of Aβ are produced, mostly ranging from 38 to 43 amino acids [2]. After its release as a monomer, Aβ, particularly in its longer forms (Aβ_42_, Aβ_43_), is very prone to self-assembly. In AD, this ultimately leads to the formation of amyloid fibrils, aggregating into senile plaques in the brain. Many studies suggest that the toxicity of Aβ does not lie in these insoluble fibrils but rather in its soluble oligomeric intermediates, as a result of their intrinsically misfolded nature and aggregation propensity that might contribute to trapping vital proteins or cause cell membrane alterations [3, 4]. Further, Aβ has been reported to have seeding properties. A particular study showed that Aβ-rich brain extracts are capable of inducing cerebral amyloidosis when inoculated into APP transgenic mice, but not into APP *knockout* mice. However, brain extracts from these previously inoculated APP *knockout* mice, that themselves do not bear amyloidosis, were still capable of inducing amyloidosis when in turn inoculated into APP transgenic mice [5]. This model of secondary transmission reveals the potential of Aβ to stably persist in the brain and also to retain pathogenic activity by acting as seeds when in the presence of host Aβ that can be propagated. Hence, it is believed that the stability of Aβ, and particularly Aβ oligomers, gives them persistent and aggravating pathological properties for the formation of amyloid deposits [6].

Aβ assembly is thought to rely on a process of nucleated polymerization [7–9], involving a nucleation phase during which unfolded or partially folded Aβ monomers self-associate to form an oligomeric nucleus. This will give rise to protofibrils through further elongation, and eventually to mature amyloid fibrils. Many Aβ assemblies have been identified as important intermediates of this pathway converting Aβ monomers into amyloid fibrils. The so-called on-pathway assemblies range from low-molecular-weight oligomers including dimers, trimers, tetramers and pentamers, to midrange-molecular-weight oligomers including hexamers, nonamers and dodecamers [10–14]. In addition, globular non-fibril forming Aβ aggregates have also been identified and labelled as off-pathway assemblies [15]. These include annulus and amylospheroid structures as well as dodecameric Aβ-derived diffusible ligands (ADDLs).

Many oligomeric structures seem to play an important role in Aβ assembly and to have deleterious effects that could explain Aβ-related toxicity [10, 16–21]. Among these assemblies, Aβ hexamers currently gain increasing interest. Indeed, they have recently been identified, together with pentamers, as the smallest oligomeric species that Aβ_42_ forms in solution [22]. Other studies based on mass spectrometry reported these assemblies as the previously mentioned nucleus (also termed “paranucleus”) that can serve as a building block for the elongation step of Aβ assembly [15, 23, 24]. This idea is supported by the fact that at least four other Aβ species are comprised of multiples of this basic hexameric unit [15, 17, 25, 26].

Altogether, these findings prompt a more in-depth understanding of the role of the different Aβ assemblies in fibrillation and deposition, but also in unravelling their intrinsic toxic properties, particularly in the case of the potentially nucleating hexamers discussed above. In this context, we previously reported the presence of ^~^28kDa oligomeric assemblies in cell lysates and culture media of CHO (Chinese hamster ovary) cells expressing amyloidogenic fragments of human APP [27]. The data reported herein demonstrate that these assemblies likely correspond to Aβ_42_ hexamers. Importantly, we identified hexameric Aβ assemblies across several cell lines, as well as in a well-established mouse model of amyloid pathology (5xFAD [28]) and in the cerebrospinal fluid (CSF) of cognitively impaired patients. We isolated hexameric Aβ from CHO cells and assessed its cytotoxic potential *in vitro* on primary neurons, as well as its potential to drive amyloid deposition *in vivo* following intracerebral stereotaxic injection. Very recent biochemical studies from our group revealed the ability of cell-derived hexameric Aβ to enhance the aggregation of synthetic Aβ monomers [29]. To address the pathological properties of hexameric Aβ in the brain, we used two mouse models: WT mice (C57BL/6) to measure the ability of the hexamers to form amyloid deposits in a non-pathological context, and transgenic 5xFAD mice to study their effect on the spreading of amyloid pathology. We found that cell-derived hexameric Aβ is a stable assembly that does not induce toxic effects by itself, but nucleates Aβ assembly in a pathological context where human Aβ accumulates (5xFAD).

As mentioned above, the production of Aβ relies on the cleavage of the βCTF membrane fragment of APP by the γ-secretase complex [1]. The catalytic core of γ-secretase is formed by either presenilin 1 (PS1) or presenilin 2 (PS2) [30, 31]. A recent cryo-EM study revealed a similar structure and conformational state between both presenilins (PSs), suggesting similar catalytic activities, but differences in their membrane-anchoring motifs [32]. We and others have repeatedly shown a major contribution of PS1 to Aβ production intended as substrate cleavage [33–37], rendering the exact role of PS2 in the amyloid pathology less understood, and in turn PS2 less attractive as a therapeutic target. However, more recent findings revealed the enrichment of PS2 γ-secretases in endosomal compartments, which might explain its lesser overall contribution to substrate cleavage simply by secondary encounters; substrates such as APP and its CTFs are indeed likely to be efficiently processed in cells first along trafficking through the secretory pathway and at the plasma membrane before reaching the endosomal pathway [38]. Further, PS2-dependent γ-secretases were shown to produce the intracellular pool of Aβ, in which longer forms prone to aggregation are accumulating [39]. Interestingly, extracellular vesicles (EVs), which are enclosed lipid membrane entities stemmed from cells circulating in biological fluids, are increasingly referred to as mediators of AD pathogenesis [40]. EVs could thus be involved in the extracellular release of specific Aβ assemblies and resulting seeding.

Together, these observations indicate that understanding the respective contribution of PS1- and PS2-dependent γ-secretases and related cellular pathways in the production of Aβ species that underpin the pathological processes at stake is of critical importance. In this context, we developed a model of stable human neuroblastoma-derived SH-SY5Y cell lines *knockdown* for each of the two PSs using the CRISPR-Cas9 homology-directed repair (HDR) technique [41, 42]. We assessed the profile of Aβ production in these cells in an effort to understand the contribution of PS1- and PS2-dependent γ-secretases in the formation of the Aβ hexamers discussed above. Our findings reveal a potentially important role of PS2 in the extracellular release of hexameric Aβ inside EVs, in agreement with the recent findings re-appraising the role of PS2 in the amyloid pathology.

## Materials and methods

### Chemicals and reagents

Reagents used for Western blotting – Pierce BCA protein assay kit, SeeBlue™ Plus2 pre-stained standard, NuPAGE™ 4-12% Bis-Tris protein gels, NuPAGE™ MES SDS Running Buffer (20X), NuPAGE™ Transfer Buffer (20X), nitrocellulose 0.1μm membranes and GE Healthcare ECL Amersham™ Hyperfilm™ – were purchased from ThermoFisher (Waltham, MA, USA). Western Lightning^®^ Plus-ECL was from Perkin Elmer (Waltham, MA, USA). Complete™ protease inhibitor cocktail was from Roche (Basel, Switzerland). Primary antibodies targeting human Aβ; anti-Aβ clone W0-2 (MABN10), anti-Aβ_40_ and anti-Aβ_42_ were from Merck (Kenilworth, MJ, USA); anti-Aβ clone 6E10 (803001) was from BioLegend (San Diego, CA, USA). Primary antibody directed against the C-terminal part of APP (APP-C-ter) (A8717) was purchased from Sigma-Aldrich (St-Louis, MO, USA). Anti-PS1 (D39D1) and anti-PS2 (D30G3) antibodies were from Cell Signalling (Danvers, MA, USA). Anti-α-tubulin primary antibody as well as secondary antibodies coupled to horseradish peroxidase (HRP) were obtained from Sigma-Aldrich. AlexaFluor-labelled secondary antibodies were obtained from ThermoFisher. Thioflavin T (ThT) amyloid stain was obtained from Sigma-Aldrich. Mowiol^®^ 4-88 used for mounting medium was purchased from Merck (Kenilworth, MJ, USA). Cell culture reagents – Ham’s-F12, DMEM-F12, DMEM and Neurobasal^®^ growth media, penicillin-streptomycin (p-s) cocktail, Lipo2000^®^ transfection reagent, Opti-MEM^®^, HBSS, glutamine and B-27^®^ – were all purchased from ThermoFisher. Fetal bovine serum (FBS) was from VWR (Radnor, PA, USA). GELFrEE™ 8100 12% Tris Acetate cartridge kits were purchased from Expedeon (Heidelberg, Germany). ReadyProbes^®^ cell viability assay kit was from ThermoFisher. Reagents used for plate-based immuno-Europium assay were ELISA strip plate (F8, high-binding 771261) from Greiner Bio-One, reagent diluent-2 10x (DY995) from R&D systems. Anti-CD9 primary antibody (MAB1880) was from R&D systems, anti-CD81 (TAPA-1, 349502) from Biolegend, anti-CD63 (MCA2142) from Serotec Bio-Rad and anti-GM130 (610823) from BD transduction. The anti-mouse IgG-biotin (NEF8232001EA), Europium-labeled streptavidin (1244-360), Delfia wash concentrate 25x (4010-0010), Delfia assay buffer (1244-111) and Delfia enhancement solution (1244-105) were all from Perkin-Elmer.

### DNA constructs

The pSVK3-empty (EP), -C42 and -C99 vectors used for expression in rodent cell lines (CHO, MEF) were described previously [27, 43]. C42 and C99 are composed of the human APP signal peptide fused to the Aβ 42 and βCTF sequences, respectively. For expression in human cell lines (HEK293, SH-SY5Y), the C99 construct in a pCDNA3.1 plasmid was kindly provided by R. Pardossi-Piquard (University of Sophia Antipolis, Nice, France). The pCDNA3.1 plasmid bearing the C99-GVP construct used in reporter gene assays was a gift from H. Karlström (Karolinska Institute, Stockholm, Sweden). The associated Gal4RE-*Firefly* luciferase reporter gene (pG5E1B-luc) and *Renilla* luciferase reporter vector (pRL-TK) have been described previously [37, 44].

### Cell lines culture and transfection

Chinese hamster ovary (CHO) cell lines were grown in Ham’s F12 medium. Human neuroblastoma SH-SY5Y and mouse embryonic fibroblasts (MEF) cell lines were grown in DMEM-F12. Human embryonic kidney (HEK293) cell lines were grown in DMEM. All media were supplemented with 10% of heat-inactivated FBS and 1% of p-s solution (100units/ml final). All cell cultures were maintained at 37°C in a humidified atmosphere and 5% CO_2_.

For transient transfection, cells were seeded 24h before transfection at a density of 40.000cells/cm^2^. Transfection mixes containing desired DNA and Lipo2000^®^ were prepared in Opti-MEM^®^ and pre-incubated for 15min at room temperature (rt) to allow for liposomal complex formation. One day after transfection, medium was changed to fresh FBS-free culture medium and incubated for another 24h. Cell lysates and culture media were harvested 48h after transfection for analysis.

### Western blotting

Cells were rinsed and scraped in phosphate-buffered saline (PBS) and centrifuged for 5min at 7.000 x *g*. Pellets were sonicated in lysis buffer (125mM Tris pH 6.8, 20% glycerol, 4% SDS) with Complete™ protease inhibitor cocktail. Protein concentration was determined using the BCA protein assay kit (Pierce, Rockford, IL, USA). Fifteen (for detection with anti-PS1 or PS2 primary antibodies) or forty (for detection with anti-Aβ or APP-C-ter primary antibodies) micrograms of protein were heated for 10min at 70°C in loading buffer (lysis buffer containing 50mM DTT and NuPAGE™ LDS sample buffer (ThermoFisher)). Samples were loaded and separated by SDS-PAGE electrophoresis on Nupage™ 4-12% Bis-Tris gels with MES SDS Running buffer, using SeeBlue™ Plus2 pre-stained as a protein standard. Proteins were then transferred for 2h at 30V with NuPAGE™ transfer buffer onto 0.1μm nitrocellulose membranes. After blocking (5% non-fat milk in PBS-Tween 0.1%), membranes were incubated overnight at 4°C with the primary antibodies, then washed, and incubated with the secondary antibodies coupled to HRP for 1h prior to ECL detection. Primary antibodies were used as follows: anti-human Aβ clone W0-2 (1:1.500), anti-human Aβ clone 6E10 (1:1.500), anti-APP-C-ter (1:2.000), anti-PS1 (1:1.000), anti-PS2 (1:1.000), anti-β-actin (1:3.000) or anti-α-tubulin (1:3.000). Secondary antibodies were used as follows: HRP-coupled anti-mouse IgG (1:10.000) or anti-rabbit IgG (1:10.000).

### GELFrEE isolation of cell-derived hexameric Aβ

Forty-eight hours after transfection with either pSVK3-EP, -C42 or -C99, the culture media of CHO cells was collected, lyophilized, re-suspended in ultrapure water and pre-cleared with recombinant protein A sepharose (GE Healthcare, Chicago, IL, USA). Immunoprecipitation of Aβ species from the media was performed with the monoclonal anti-Aβ clone W0-2 (MABN10) antibody. Samples were separated through a Gel Eluted Liquid Fraction Entrapment Electrophoresis (GELFrEE™) 8100 system to allow the collection of the desired kDa range of proteins directly in liquid fraction. The following method was used; step 1: 60min at 50V, step 2: 6min at 70V, step 3: 13min at 85V and step 4: 38min at 85V. Fractions 1, 2 and 3 (Fig1) were collected in the system running buffer (1X buffer; 1% HEPES, 0.01% EDTA, 0.1% SDS and 0.1% Tris) at the end of step 2, 3 and 4 respectively. Samples were aliquoted and kept at −20°C until use.

**Fig1.**
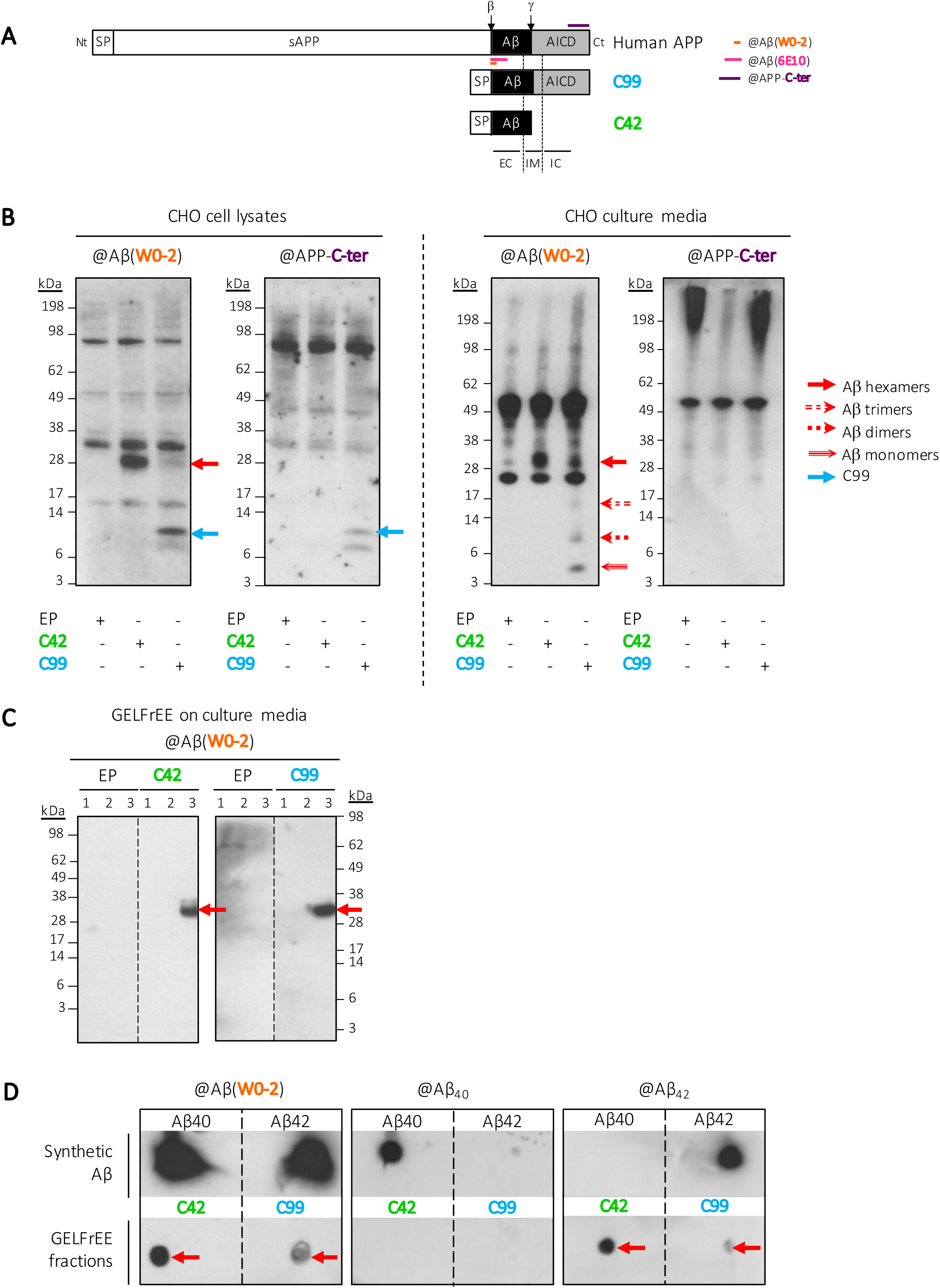
Detection of hexameric Aβ_42_ in CHO cells expressing human APP metabolites. **A.** Full-length APP is cleaved by β-secretase at the β site, located at the N-terminus of Aβ, to produce a 99-amino acid long membrane-bound fragment called βCTF (encompassing Aβ and AICD). The construct referred to as C99 here is composed of the signal peptide (SP) of APP fused to the sequence of the βCTF fragment. This fragment is then cleaved by γ-secretase at the γ site to release Aβ. The C42 construct is composed of the SP of APP and the Aβ_42_ sequence. The epitopes of the primary antibodies directed against human Aβ (clones W0-2 and 6E10) and the C-terminal region of APP (APP-C-ter) used in this study are indicated on the scheme. Nt: N-terminus. Ct: C-terminus. sAPP=soluble APP. AICD=APP intracellular domain, EC=extracellular, IM=intramembrane, IC=intracellular. **B.** Detection of an assembly of ^~^28kDa by Western blotting in CHO cell lysates and culture media following expression of either C42 or C99. This assembly is recognized by the W0-2 antibody, but not by the APP-C-ter, suggesting it emerges by assembly of Aβ. In the media of C99-expressing cells, intermediate assemblies are also observed; monomers, dimers and trimers. EP=empty plasmid. **C.** Isolation of cell-derived Aβ. The media of CHO cells expressing either EP, C42 or C99 were immunoprecipitated and separated using the GELFrEE technique. We optimized a method to collect the ^~^28kDa Aβ assembly as an isolated liquid fraction. Dashed lines indicate that proteins were run on the same gel, but lanes are not contiguous. **D.** Dot blotting of the isolated ^~^28kDa assemblies revealed they are composed of Aβ_42_. Synthetic preparations of monomeric Aβ_40_ and Aβ_42_ were used as positive controls. Combined with the observed size, we identify the assemblies of interest as Aβ_42_ hexamers. Dashed lines indicate that proteins were loaded on the same membrane, but image was readjusted.

### Dot blotting

5μl of isolated hexameric Aβ (150μM for isoform characterization or 15μM for fractions evaluation prior to intracerebral injection) and 5μl of 50μM synthetic monomeric Aβ_40_ or Aβ_42_ were spotted onto 0.1μm nitrocellulose membranes and allowed to dry. Another 5μl of sample were then spotted twice on top and dried. The membranes were boiled twice in PBS for 3min, then blocked with 5% non-fat milk in PBS-Tween 0.1% as for Western blotting. Membranes were then washed and incubated with primary and secondary antibodies prior to ECL detection as described above. Primary antibodies dilutions were used as follows: anti-human Aβ clone W0-2 (1:1.500), anti-Aβ_40_ (1:1.000), anti-Aβ_42_ (1:1.000). Secondary antibodies were used as described for Western blotting.

### Animal models

Transgenic 5xFAD mice (Tg6799) harboring human APP and PS1 transgenes were originally obtained from the Jackson Laboratory: B6SJL-Tg(APPSwFlLon,PSEN1*M146L*L286V)6799Vas/Mmjax (34840-JAX). Colonies of 5xFAD and non-transgenic (wild-type, WT) mice were generated from breeding pairs kindly provided by Pr. Jean-Pierre Brion (ULB, Brussels, Belgium). All mice were kept in the original C57BL/6 background strain. Animals were housed with a 12h light/dark cycle and were given *ad libitum* access to food and water. All experiments conducted on animals were performed in compliance with protocols approved by the UCLouvain Ethical Committee for Animal Welfare (reference 2018/UCL/MD/011).

### Protein extraction from mouse brain tissues

WT and 5xFAD mice were euthanized by cervical dislocation or using CO_2_ and brains were quickly removed. The hippocampus as well as a portion of temporal cortex were immediately dissected on ice. Brain tissues were then homogenized by sonication in ice-cold lysis buffer (150mM NaCl, 20mM Tris, 1% NP40, 10% glycerol) with Complete™ protease inhibitor cocktail until homogenous. Samples were stored at −80°C until use. Protein concentration was determined using the BCA protein assay kit prior to analysis.

### Cerebrospinal fluid collection

Cerebrospinal fluid (CSF) was collected by lumbar puncture from AD patients and symptomatic controls undergoing diagnostic work-up at the Cliniques Universitaires Saint-Luc (UCL, Brussels, Belgium), following the international guidelines for CSF biomarker research [45]. Collected samples were directly frozen at −80°C until analysis and were always manipulated on ice during Western blotting and ECLIA experiments. Included patients signed an internal regulatory document, stating that residual samples used for diagnostic procedures can be used for retrospective academic studies, without any additional informed consent (ethics committee approval: 2007/10SEP/233). AD patients participated in a specific study referenced UCL-2016-121 (Eudra-CT: 2018-003473-94). In total, CSF samples from eight subjects were retrospectively monitored in this study (see Additional File 2).

### Electro-chemiluminescence immunoassay (ECLIA) for monomeric Aβ quantification

Aβ monomeric peptides were quantified in CSF samples as well as in the culture media of SH-SY5Y cells using the Aβ multiplex ECLIA assay (Meso Scale Discovery, Gaithersburg, MD, USA) as previously described [46]. Aβ was quantified according to the manufacturer’s instructions with the human Aβ specific 6E10 multiplex assay. For SH-SY5Y, cells were conditioned in FBS-free medium for 24h and cell medium was collected, lyophilized and re-suspended in ultrapure water.

### Primary neuronal cultures

Primary cultures of neurons were performed on E17 mouse embryos as described previously [47]. Briefly, cortices and hippocampi were isolated by dissection on ice-cold HBSS and meninges were removed. Tissues were then dissociated by pipetting up and down 15 times with a glass pipette in HBSS-5mM glucose medium. Dissociation was repeated 10 times with a flame-narrowed glass pipette and allowed to sediment for 5min. Supernatant containing isolated neurons was then settled on 4ml FBS and centrifuged at 1.000 x *g* for 10min. The pellet was resuspended in Neurobasal^®^ medium enriched with 1mM L-glutamine and 2% v/v B-27^®^ supplement medium. Cells were plated at a density of 100.000cells/cm^2^ in 12w plates pre-coated with poly-L-lysine (Sigma-Aldrich). Cultures were maintained at 37°C and 5% CO_2_ in a humidified atmosphere. Half of the medium was renewed every 2 days and neurons were cultured for 8 days *in vitro* (DIV8) before being used for cell viability experiments.

### Cell viability assay (ReadyProbes^®^)

To assess the toxicity of cell-derived hexameric Aβ *in vitro*, primary neuronal cultures performed from WT and 5xFAD mouse embryos were incubated (at DIV7) with 1 or 5μM of either cell-derived hexameric Aβ (C42 fraction) or control (EP fraction). At DIV8, 2drops/ml of each reagent of the ReadyProbes^®^ assay were added to cells: NucBlue^®^ Live reagent for the staining of all nuclei and NucGreen^®^ Dead reagent for the nuclei of cells with compromised plasma membrane integrity. Staining were detected with standards DAPI and FITC/GFP filters respectively, at an EVOS^®^ FL Auto fluorescence microscope. Quantification was performed by counting dead vs total cells on ImageJ.

### Intracerebral stereotaxic surgery

Stereotaxic injections of isolated Aβ hexamers were performed in WT and 5xFAD mice to analyze the effect of cell-derived β-amyloid hexamers on the development of Aβ pathology *in vivo*. 2-month-old mice were deeply anesthetized by intra-peritoneal injection of a mixture of ketamine (Ketamin^®^) (10mg/kg) and medetomidine (Domitor^®^) (0.5mg/kg) and placed in a stereotaxic apparatus (Kopf^®^ Instruments). 2μl of 15μM cell-derived Aβ hexamers (C42 fraction) or control (EP fraction) were injected using a 10μl Hamilton syringe and an automated pump (RWD^®^) in the hippocampus (A/P −1.94; L +-2.17; D/V −1.96; mm relative to bregma). 30 days after stereotaxic injection, mice were transcardially perfused with PBS and brains were post-fixed in 4% paraformaldehyde for 24h at 4°C.

### Immunohistofluorescence

For immunohistological analysis, free-floating coronal sections (50μm) were generated from agarose-embedded fixed brains using a vibrating HM650V microtome and were preserved in PBS-sodium azide 0.02% at 4°C. Prior to immunomarking, sections were washed in PBS and subsequently blocked and permeabilized with PBS-BSA 3%-TritonX100 0.5% for 1h at rt. Sections were then incubated with antihuman Aβ clone W0-2 (MABN10, 1:100) overnight at 4°C as a marker for Aβ. After three PBS washes and incubation with AlexaFluor647-coupled secondary antibody (1:500) for 1h at rt, slices were finally washed three times with PBS and mounted on SuperFrost^®^ slides. Slides were then incubated with ThT (0.1mg/ml in ethanol 50%) for 15min at rt as a marker for fibrillar deposits. After three washes with ethanol 80% and a final wash with ultrapure water, coverslips were mounted with Mowiol^®^ 4-88-glycerol. W0-2 and ThT staining were detected with FITC/Cy5 and GFP respectively at an EVOS^®^ FL Auto fluorescence microscope. Counting of double-positive dots was performed on ImageJ.

### Generation of SH-SY5Y PS1 and PS2 deficient cells by CRISPR-Cas9

To generate PS1 and PS2 deficient cells, kits each containing 2 guide RNA (gRNA) vectors that target human *PSEN1* or *PSEN2* genes, a GFP-puromycin or RFP-blasticidin donor vector respectively, and a scramble control were obtained from Origene (CAT#: KN216443 and KN202921RB). Target sequences were flanked with specific homology sequences for the stable integration of donor sequences, based on the homology-directed repair technique [41, 42]. SH-SY5Y cells were transfected using Lipo2000^®^ according to the manufacturer’s instructions (*cf*. Cell culture and transfection section). Two days after transfection, cells were FACS-sorted for GFP+ (PS1) or RFP+ (PS2) cells and seeded in 24w plates. Following a few days for cell-growth, a second selection was performed using puromycin (PS1) or blasticidin (PS2) at a concentration of 15μg/ml or 30μg/ml, respectively. Cells were allowed to grow again and split twice before sub-cloning in 96wells. Clonal populations were then assessed for PSs protein expression by Western blotting. PS1 and PS2 clonal populations were selected for following experiments on account of the highest gene-extinction efficiency. Puromycin (2.5μg/ml) and blasticidin (7.5μg/ml) were used for the maintenance of PS1 and PS2 deficient cells, respectively.

### Dual luciferase assay

SH-SY5Y cells were co-transfected with Lipo2000^®^ (*cf*. Cell culture and transfection section) in a 1:1:1 ratio with pG5E1B-luc, pRL-TK and either a pCDNA3.1-empty plasmid (EP) or the pCDNA3.1-C99-GVP, bearing a tagged C99 to quantify the release of AICD. The system setup was previously described [37, 48]. Forty-eight hours after transfection, cells were rinsed with PBS and incubated with the reporter lysis buffer (Promega, Madison, WI, USA) for 15min at rt. *Firefly* and *Renilla* luciferase activities were measured using the Dual-Glo^®^ Luciferase Assay System (Promega, Madison, WI, USA) on a Sirius single tube luminometer (Berthold, Bad Wildbad, Germany). Luciferase activity corrected for transfection efficiencies was calculated as the *Firefly/Renilla* ratio.

### Extracellular vesicles isolation

Ultracentrifugation was performed on media from SH-SY5Y scramble and *knockdown* cells for the isolation of vesicular entities, as previously described [49]. Briefly, culture medium was collected and underwent several centrifugation steps (all at 4°C); 300 x *g* for 10min for the elimination of living cells, 1.000 x *g* for 10min to discard dead cells, 10.000 x *g* for 30min for the removal of cellular debris, and finally 100.000 x *g* for 1h, to collect extracellular vesicles (EVs) as a pellet and soluble proteins as supernatant. Soluble proteins were precipitated by incubation with 10% v:v trichloroacetic acid (TCA) for 30min on ice. Both EVs and soluble proteins fractions were resuspended in 500μl of PBS for nanoparticle tracking analysis (NTA) and plate-based immuno-Europium-assay. For Western blotting, both fractions were sonicated in lysis buffer (125mM Tris pH 6.8, 20% glycerol, 4% SDS) with Complete™ protease inhibitor cocktail. Protein concentration was determined using the BCA protein assay kit.

### Nanoparticle tracking analysis (NTA)

EVs were counted in each fraction by the Zetaview (ParticleMetrix,GmbH), which captures Brownian motion through a laser scattering microscope combined with a video camera to obtain size distribution (50-1000nm) and concentration. Samples were diluted 1:50 in PBS to reach 50-200 particles/frame, corresponding to ^~^2.107-1.108 particles/ml. Sensitivity was set to 65 and camera shutter to 100 in order to detect less than 3 particles/frame when BPS alone was injected, to assess background signal. Measurements were averaged from particles counted in 11 different positions for 2 repeated cycles with camera at medium resolution mode.

### Plate-based Europium-immunoassay

50μl of EVs and soluble fractions were bound to protein-binding ELISA plates. After overnight incubation at 4°C, the rest of the experiment was performed at rt by shaking on a tilting shaker at 30rpm. The plate was washed with Delfia buffer (diluted to 1x in PBS; Delfia-W), then blocked with reagent diluent-2 (diluted to 1% BSA in PBS) for 90min. The bound material was labelled with primary antibodies against CD9, CD81, CD63 and GM130 (1μg/ml in reagent diluent-2 diluted to 0.1% BSA in PBS) for 90min. After three Delfia-W washes, goat anti-mouse biotinylated antibody (1:2.500 in reagent diluent-2 diluted to 0.1% BSA in PBS) was added for 60min. After three Delfia-W washes, Europium-conjugated streptavidin (diluted to 1:1.000 in Delfia assay buffer assay) was added for 45min. After six final Delfia-W washes, Delfia enhancement solution was incubated for 15min before measurement using time-resolved fluorometry with exc/em 340/615nm, flash energy/light exposure high/medium and integration lag/counting time 400/400μs (Victor X4 multilabel plate reader, PerkinElmer).

### Statistical analyses

The number of experiments (N) and the number of samples per condition in each experiment (n) are indicated in figure legends. All statistical analyses were performed using the GraphPad Prism 8 software (GraphPad Software, La Jolla, CA, USA). Gaussian distribution was assessed using the Shapiro-Wilk test. A parametric test was applied if the data followed normal distribution. Otherwise, non-parametric tests were used. When tested groups were each expressed as a fold-change of their corresponding control, the value of the control was set as the hypothetical value for the use of parametric one-sample *t* test or non-parametric one-sample Wilcoxon single-ranked test. When a correlation between two variables was assessed, Pearson’s R correlation coefficient was calculated. When two groups were compared, parametric *t* test with Welch’s correction or non-parametric Mann-Whitney test were used. When more than two groups were compared, parametric ANOVA with indicated *post hoc* tests or non-parametric Kruskal-Wallis were used. Significance is indicated as: ns=non-significant, *=p<0.05, **=p<0.01, ***=p<0.001. Actual p-values of each test are indicated in the corresponding figure legend.

## Results

### Identification of cell-derived hexameric Aβ_42_ in CHO cells expressing human APP metabolites

Considering the emerging body of evidence pointing to the pathological properties of oligomeric Aβ assemblies, we first assessed the profile of Aβ production in a widely used cell model; the CHO (Chinese hamster ovary) cell line. We transiently transfected these cells with vectors expressing the human sequences of either Aβ_42_ or βCTF, each fused to the signal peptide of the full-length APP protein (Fig1A) to ensure a proper cellular trafficking of the expressed fragments. These constructs will be referred to as C42 and C99 respectively. The cell-derived Aβ assemblies were analyzed by Western blotting with several monoclonal antibodies targeting the human Aβ sequence (W0-2 and 6E10 clones) or the APP C-terminal region (APP-C-ter). Strikingly, we reproducibly detected a band at ^~^28kDa in both C42 and C99 conditions with Aβ-specific antibodies (W0-2 clone in Fig1B and 6E10 clone in Additional File 1). This assembly is not recognized by the APP-C-ter specific antibody (Fig1B), supporting the idea that it is formed by assembly of the Aβ fragment, and likely corresponds to an Aβ_42_ hexamer. Importantly, we noticed a key difference between C42 and C99 expressing cells; when C42 is expressed, only the hexameric Aβ assembly is detected in both cell lysates and media, while in the C99 condition, which requires processing by γ-secretase to release Aβ, we detect several intermediate Aβ assemblies; monomers, dimers, trimers and hexamers (Fig1B). This suggests that the Aβ, whether directly expressed or released by processing in the cellular context, will either readily or progressively assemble into the identified hexameric assembly. In addition, as the presence of these low-molecular-weight assemblies is only observed in the media of C99 expressing cells and not inside the cells, the ranges of Aβ assemblies may be produced differentially or aggregate with different kinetics depending on the extra- or intracellular context in which they accumulate.

For a further characterization of the Aβ assembly identified in Fig1B, we used a Gel Elution Liquid Fraction Entrapment Electrophoresis technique (GELFrEE™ 8100) to isolate the cell-derived Aβ hexamers from W0-2-immunoprecipitated media of CHO cells expressing C42 or C99 (Fig1C). Dot blotting with primary antibodies directed against the free C-terminal end of the two major Aβ isoforms (Aβ_40_, Aβ_42_) was performed on the isolated ^~^28kDa fraction and identified it as an Aβ_42_ assembly (Fig1D). Synthetic preparations of monomeric Aβ_40_ and Aβ_42_ were used as positive controls. Altogether, these results converge to the identification of the assembly of interest as cell-derived hexameric Aβ_42_.

### Formation of hexameric Aβ across a wide range of models

As Aβ self-assembly strongly depends on the context of its release, we sought to determine whether the assembly of interest was produced particularly by CHO cells or commonly across other models. Using the same procedure as described above, we assessed the Aβ profile in transiently transfected mouse embryonic fibroblasts (MEF) (Fig2A) as well as two human immortalized models – human embryonic kidney (HEK293) cells (Fig2B) and neuroblastoma-derived SH-SY5Y cells (Fig2C). Importantly, the ^~^28kDa assembly was consistently detected with the W0-2 Aβ specific antibody and not by the APP-C-ter targeted antibody in all the tested models (Fig2). This indicates that Aβ hexamers can readily form in a cellular context and are not restricted to one cell-type, fostering its interest as a primary nucleus for Aβ toxicity and amyloid pathology.

**Fig2.**
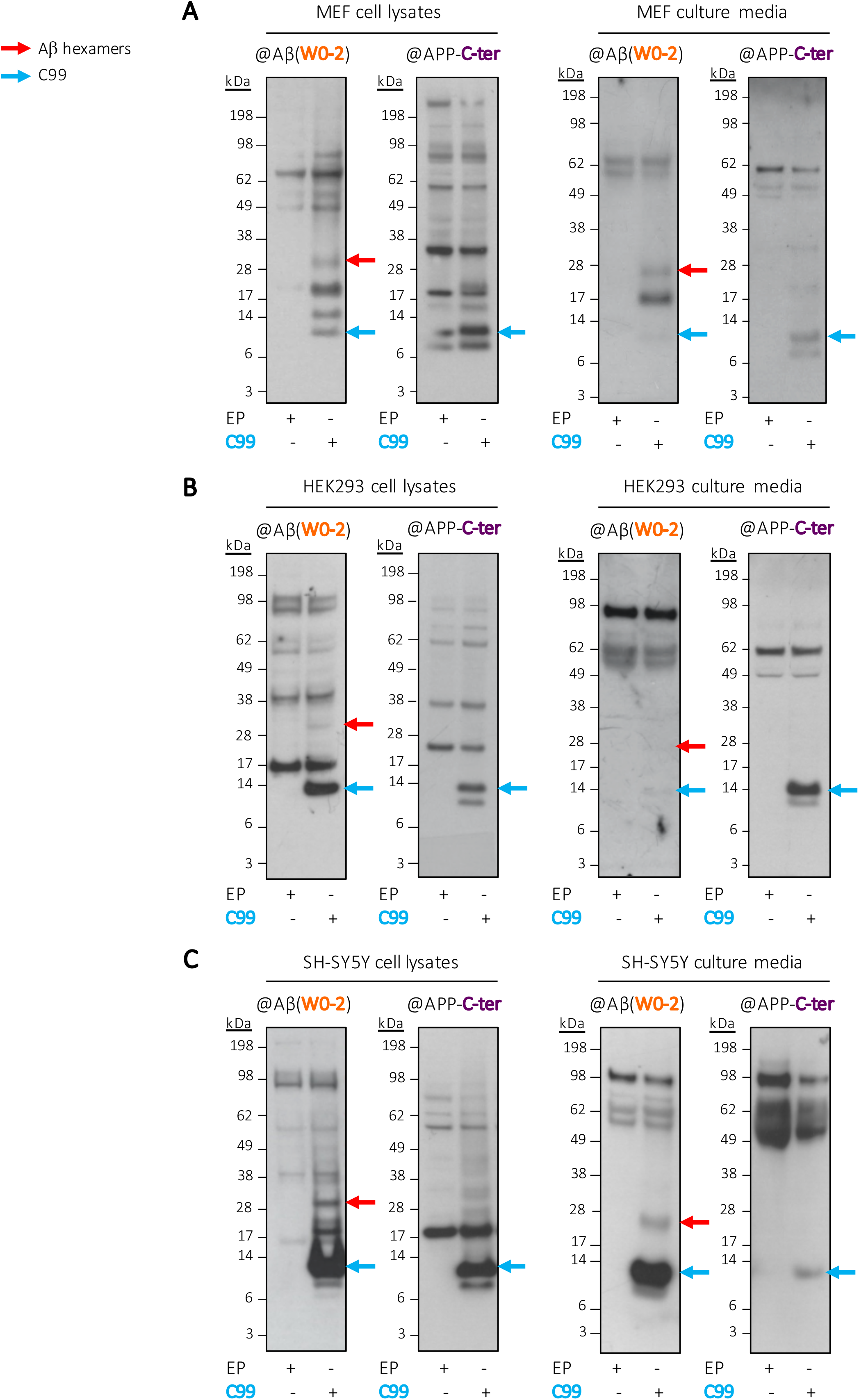
Commonality of hexameric Aβ production in several cell lines. Cell lysates and media from human neuroblastoma SH-SY5Y (in **A.**) and embryonic HEK293 (in **B.**) cells, as well as from murine MEF fibroblasts (in **C.**) expressing C99 all revealed the presence of a ^~^28kDa band recognized by the human Aβ specific W0-2 antibody, and not by the anti-APP-C-ter. EP= empty plasmid.

We next investigated the presence of this Aβ assembly in mice bearing familial AD (FAD) mutations. We readily detected the ^~^28kDa Aβ assembly (Fig3A) in brain extracts of 5xFAD mice [28]. Interestingly, the intensity of hexameric Aβ increases with age. More importantly, the detection of the hexameric assembly precedes that of fibrils, which are recognized as the major indicator of the development of amyloid deposits in the 5xFAD model [28, 50]. Quantitative analysis of hexamers and fibrils relative to human APP expressed in mice brains confirms the appearance of hexameric Aβ as an early event (Fig3B). To note, the assembly of interest accumulates first in the hippocampus of the mice, as early as 2-month-old, while its increase in cortical regions peaks at 3 to 6-month of age. This is in line with the staging of amyloid pathology observed in AD and again suggests hexameric Aβ might serve as an early indicator of pathology development, and possibly as a nucleus for further amyloid deposition.

**Fig3.**
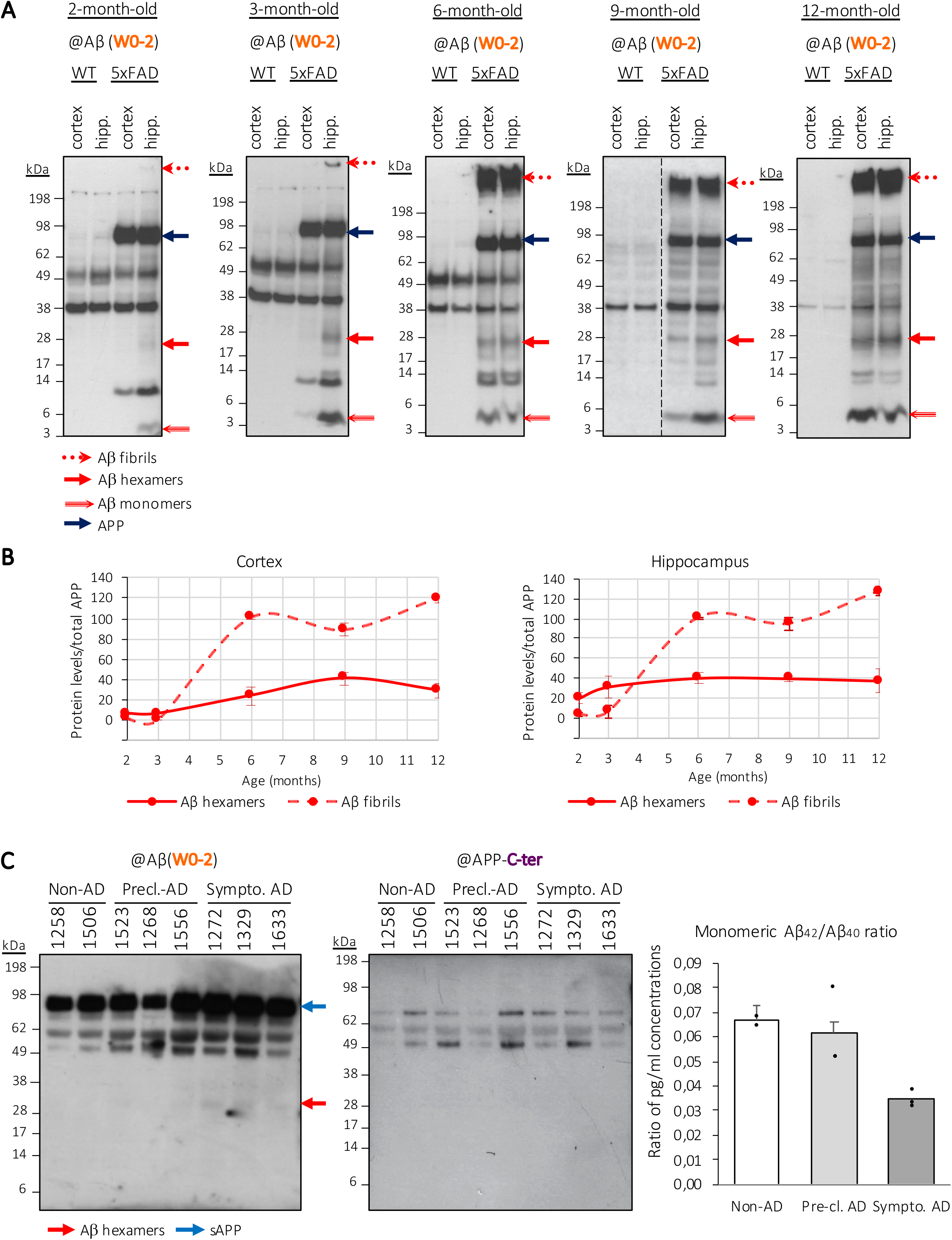
Identification of hexameric Aβ as an assembly involved in the context of AD. **A.** Detection of Aβ assemblies in brain samples from an amyloid mouse model (5xFAD). Cortices and hippocampi of euthanized mice were lysed and analyzed by Western blotting. Aβ fibrils appear stuck in the wells and hexameric Aβ assemblies are detected at ^~^28kDa. To note, Aβ monomers are also detected in all 5xFAD samples and reflect an efficient metabolism of the human APP protein expressed in these mice. Dashed lines indicate that proteins were run on the same gel, but lanes are not contiguous. Hipp.=hippocampus. **B.** The signal intensities of Aβ hexamers and Aβ fibrils were quantified relatively to the APP signal. Samples used for quantitative analysis derived from the same experiment, with Western blots processed in parallel. The displayed graphs represent the profile of Aβ assemblies as related to both the analyzed brain area (cortex, hippocampus) and the age (2, 3, 6, 9, 12 months of age) (min N=3 each). **C.** Identification of hexameric Aβ in the cerebrospinal fluid (CSF) of cognitively affected patients. Western blotting analysis was performed using the W0-2 and APP-C-ter antibodies. Dosage of monomeric Aβ_42_/Aβ_40_ by ECLIA immunoassay confirmed the correct classification of individuals, with a reduction in ratio along with AD progression.

Finally, cerebrospinal fluid (CSF) samples from cognitively affected patients (diagnosed with non-AD dementia, pre-clinical AD or symptomatic AD) were monitored with the same Western blotting approach. Only AD-related patients revealed the presence of ^~^28kDa assemblies, undeniably placing the assembly of interest in the pathological context of AD (Fig3C, left panel). More detailed information on neurological examination and PET-analyses conducted on the patients are displayed in Additional File 2. The same CSF samples were used for a quantitative analysis of monomeric A β isoforms by ECLIA immunoassay and revealed an overall reduction in the Aβ_42_/Aβ_40_ ratio in AD patients (Fig3C, right panel). This reduction correlated with the severity of AD symptoms exerted by the patients (see raw values in Table1), concordantly with previous reports [51]. On the contrary, relative quantification of hexameric Aβ levels detected by Western blotting, using soluble APP as an intrasubject control, revealed an increase in hexameric levels with the progression of AD (Table1). This suggests a correlation between the reduced proportion of monomeric Aβ_42_ and its aggregation in higher assemblies, as previously suggested [52, 53], but here particularly with the hexameric assembly. More precisely, statistical analysis revealed that 48.8% of the Aβ_42_/Aβ 40 ratio variance can be explained by the increase in hexameric Aβ formation.

**Table 1.** Inverse correlation between monomeric and hexameric Aβ in human CSF samples.

| Inclusion n° | Classification | A$\beta_{42}$/A$\beta_{40}$ | Hexameric Aβ/sAPP |
| --- | --- | --- | --- |
| 1258 | Non-AD | 0.069 | 0.079 |
| 1506 |  | 0.065 | 0.055 |
| 1523 | Pre-clinical AD | 0.052 | 0.075 |
| 1268 |  | 0.080 | 0.084 |
| 1556 |  | 0.052 | 0.092 |
| 1272 | Symptomatic AD | 0.033 | 0.140 |
| 1329 |  | 0.039 | 0.224 |
| 1633 |  | 0.032 | 0.143 |

CSF samples from eight subjects were monitored in this study: patients n°1258 and 1506 did not exert any AD-related features. Patients n°1523, 1268 and 1556 were diagnosed with pre-clinical AD and patients n°1272, 1329 and 1633 with symptomatic AD. Ratios of A $\beta_{42}$ /A $\beta_{40}$ and of hexameric A $\beta$ /soluble APP (sAPP) were obtained from ECLIA and Western blot quantifications respectively (see Fig3). Pearson's R correlation test: $R=-0.70$ ( $R\text{-squared}=0,49$ ), $p=0.05$ .

### Cell-derived hexameric Aβ causes cell viability impairment only in primary neurons expressing amyloid proteins *in vitro*

Following the detection of hexameric Aβ in a well-described amyloid mouse model and in the CSF of cognitively impaired patients, identifying it as an important factor in the development of amyloid pathology, we sought to assess whether isolated cell-derived hexameric Aβ would exert any neurotoxicity. For this, we cultured primary neurons from wild-type (WT) and transgenic (5xFAD) embryos and treated them after 7 days of differentiation *in vitro* (DIV) with hexameric Aβ. Aβ hexamers were obtained by W0-2-immunoprecipitation and GELFrEE separation of C42-expressing CHO cells media as described above (Fig1). The corresponding GELFrEE fraction of cells expressing the empty plasmid (EP) were used as a control. Two final concentrations were tested; 1μM and 5μM. 24h after treatment, cell viability was assessed using a ReadyProbes^®^ assay and a percentage of dead cells out of the total cells was quantified (Fig4A). Results show an absence of any significant cytotoxic effect at tested concentrations on primary neurons cultured from WT mice (Fig4B), even though both are above the reported neurotoxic concentrations from synthetic preparations of oligomeric Aβ [6, 54]. This suggests that the identified assembly is not cytotoxic by itself, at least in these experimental conditions. However, primary neurons cultured from 5xFAD mice, which can serve as an amyloid model *in vitro* [55–57], displayed increased cell death when exposed to 5μM of cell-derived hexameric Aβ. Importantly, this indicates that Aβ hexamers may have the ability to cause toxic effects only when there is pre-existing Aβ in the neuronal environment. This therefore implies that such a cytotoxic ability requires the intermediate seeding of other Aβ present.

**Fig4.**
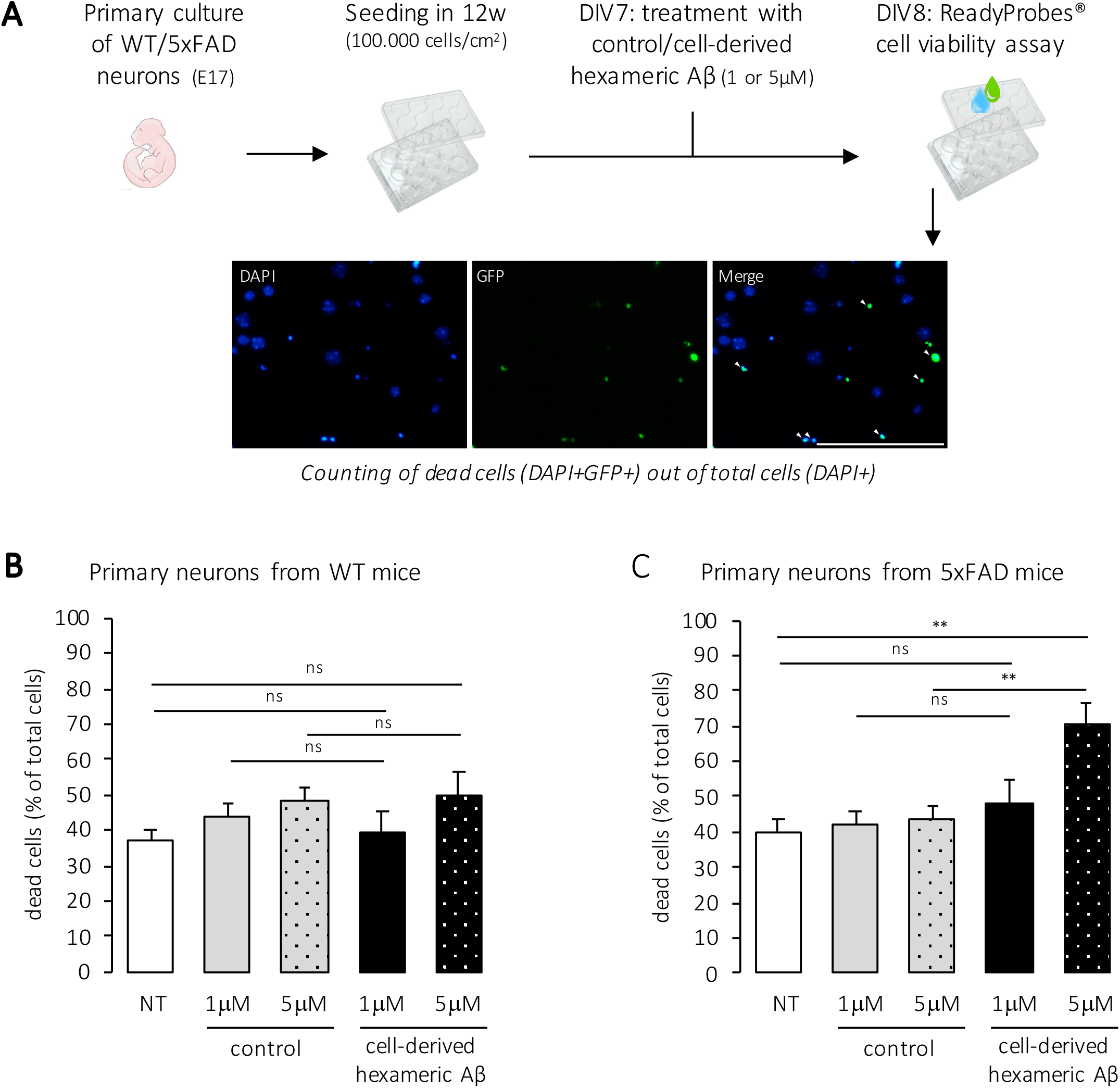
Cell-derived Aβ hexamers are only cytotoxic in primary neurons that express amyloid proteins. **A.** Experimental workflow. Primary neurons were isolated from wild-type (WT) or transgenic (5xFAD) mouse embryos at stage E17 and cultured for 8 days *in vitro* (DIV). At DIV7, cells were incubated for 24h with 1μM or 5μM of cell-derived hexameric Aβ or control, isolated from the media of C42- and EP-expressing CHO cells respectively. Cell viability was assessed using the ReadyProbes^®^ assay and fluorescent staining was captured at an EVOS^®^ FL Auto fluorescence microscope. A representative image of the assay is shown. Scale bar=50μm. **B, C.** Quantification of the proportion of dead cells compared to the total cells in WT (in **B.**) and 5xFAD cultures (in **C.**). Total number of cells counted (number of dead cells counted in brackets) was as follows in WT: n=1183(439), 1070(472), 650(314), 813(318), 797(400) and 5xFAD: n=528(212), 640(270), 1019(442), 465(224), 775(544) for NT, control (equivalent of 1μM), control (equivalent of 5μM), hexameric Aβ (1μM) and hexameric Aβ (5μM) respectively. NT=not treated. N=4 independent experiments in WT, N=3 independent experiments in 5xFAD. One-way ANOVA with Tukey’s multiple comparison test: ns=non-significant, *=p<0.05, **=p<0.01 (in WT: p=0.99 NT vs hexameric Aβ (1μM), p=0.38 NT vs hexameric Aβ (5μM), p=0.95 control (1μM) vs hexameric Aβ (1μM), p=0.97 control (5μM) vs hexameric Aβ (5μM); in 5xFAD: p=0.70 NT vs hexameric Aβ (1μM), p=0.004 NT vs hexameric Aβ (5μM), p=0.85 control (1μM) vs hexameric Aβ (1μM) and p=0.009 control (5μM) vs hexameric Aβ (5μM)).

### Cell-derived hexameric Aβ aggravates *in vivo* amyloid deposition in a transgenic mouse model

To further assess its potential to drive amyloid formation, we performed hippocampal stereotaxic injections of cell-derived hexameric Aβ in two mouse models: (i) WT mice (C57BL/6) to assess the potential of Aβ hexamers to form amyloid deposits in a previously amyloid-free context, (ii) mice developing amyloid pathology (5xFAD) to mimic a situation where the hexamers are incubated with pre-existing Aβ to serve as seeds and thus study whether they have a nucleating potential *in vivo*, driving the assembly and deposition of Aβ produced in the brain. Experimental workflow is represented in Fig5A. As for *in vitro* toxicity assays, Aβ hexamers were obtained by W0-2-immunoprecipitation and GELFrEE separation of C42-expressing CHO cells media and the corresponding fraction of EP-expressing CHO cells media was used as a control. The fraction of cell-derived hexameric Aβ was diluted from 150μM to 15μM prior to intracerebral injection. The control fraction was diluted in a similar manner. Specific detection of diluted cell-derived hexameric Aβ was confirmed by dot blotting with the W0-2 antibody (Fig5A, left panel). 2-month-old WT or 5xFAD mice were injected in the hippocampus of the left and right hemisphere with 2μl of EP (control) and C42 (cell-derived hexameric Aβ) diluted fractions, respectively. To evaluate Aβ deposition, mice were sacrificed 30 days after stereotaxic injection and brains were fixed. Coronal sections were co-stained with the human specific W0-2 antibody for Aβ and the ThT dye for fibrillar aggregates. Quantitative analysis of Aβ deposition was performed by counting double-positive dots (as indicated in Fig5A). The results showed that cell-derived hexameric Aβ does not have the ability to form fibrillar deposits by itself in a WT brain (Fig5B), but is capable of enhancing the deposition of Aβ present in the 5xFAD brain (Fig5C). In transgenic mice, the overall deposition of Aβ in the hemisphere injected with cell-derived hexameric Aβ showed a significant 1.47-fold increase when compared to the control-injected hemisphere (average (±SEM) of 32.39 (±3.49) and 47.50 (±4.74) deposits per field in control- and hexamer-injected hemispheres, respectively). Deposits were investigated in the two regions mainly affected by amyloid pathology in AD: the hippocampus and the cortical areas (Fig5C). As expected, the highest increase in Aβ deposition was observed in the hippocampal region, where stereotaxic injections were performed (2.90-fold increase). However, levels of Aβ deposits were also significantly increased by a 1.74-fold in the cortex. This suggests that the injected cell-derived hexameric Aβ is able to propagate from the hippocampal formation to associated cortical regions to promote amyloidosis.

**Fig5.**
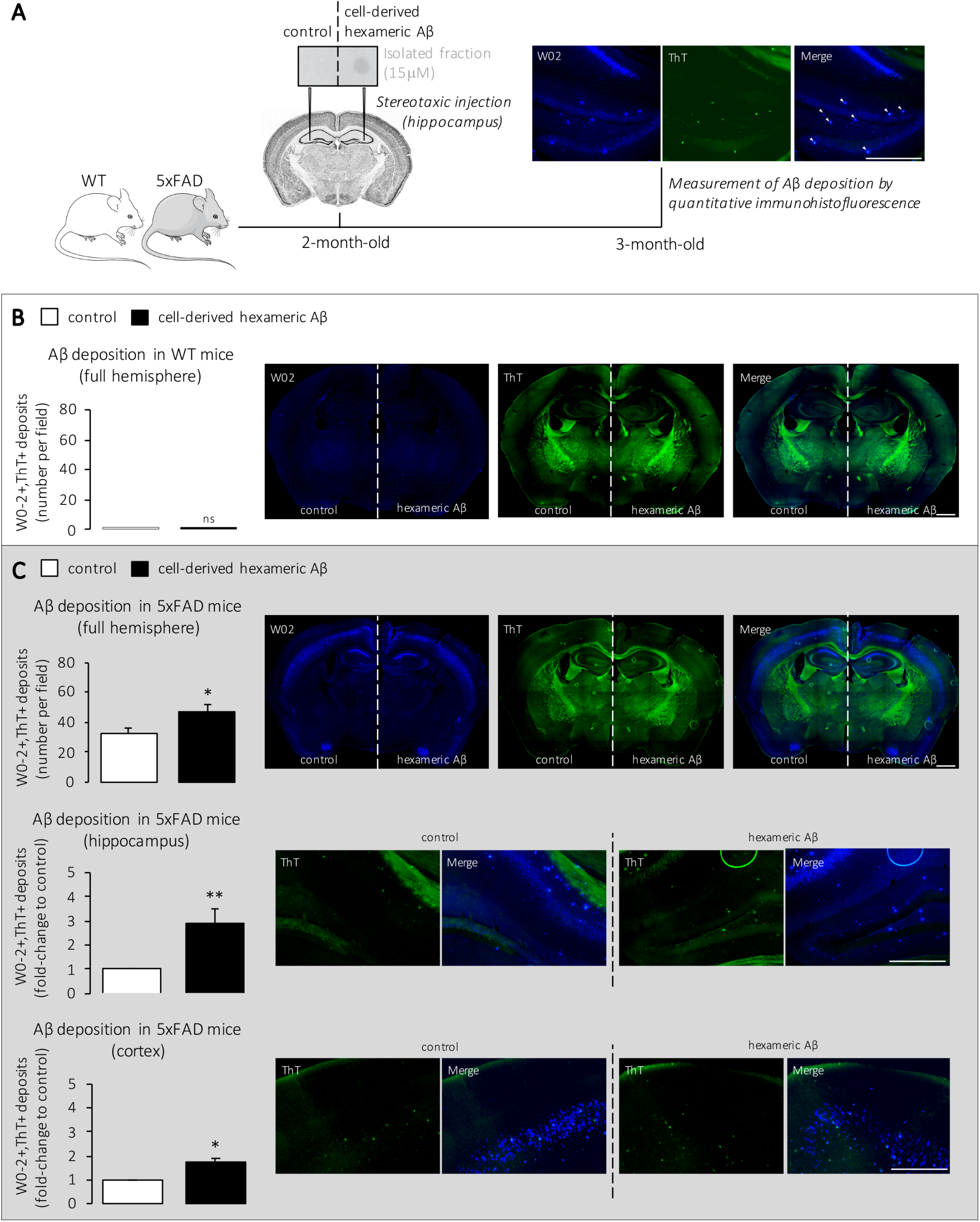
Intracerebral injection of cell-derived hexameric Aβ in WT and 5xFAD mice. **A.** Experimental workflow. 2-month-old mice were deeply anesthetized, placed in a stereotaxic apparatus and bilaterally injected with 2μl of either 15μM GELFrEE-isolated Aβ hexamers (C42) or control (EP) in the hippocampus (A/P −1.94; L +-2.17; D/V −1.96; mm relative to bregma). Both fractions were analyzed by dot blotting prior to injection. 30 days later, mice were transcardially perfused, brains were fixed and coronally sectioned (50μm) using a vibrating HM650V microtome. Immunostaining was performed on free-floating sections using the anti-human Aβ (W0-2) antibody as a marker for Aβ and the Thioflavin T (ThT) dye as a marker for fibrillar deposits. W0-2 and ThT staining were detected with FITC/Cy5 and GFP filters respectively. Right panel displays an example of double-positive counting. Scale bar=_400_μm. **B, C**. Quantification of fibrillar deposits in full hemispheres of WT (in **B.**) and 5xFAD brains (in **C.** upper panel, scale bar=1000μm) injected with control vs hexameric Aβ. n=32 slides from N=8 mice for both WT and 5xFAD. Mann-Whitney test: ns=non-significant, *=p<0.05 (in WT: p>0.99 control vs hexameric Aβ; in 5xFAD: p=0.04 control vs hexameric Aβ). For transgenic mice, deposits were also classified according to the two most affected brain regions, hippocampus and cortex, as a function of the control-injected hemisphere (in **C.** middle and lower panel, scale bar=400μm). A 2.90-fold and a 1.74-fold increase were observed in the hippocampus and cortex respectively. One-sample Wilcoxon signed-rank test with hypothetical value set at 1: *=p<0.05, **=p<0.01 (in 5xFAD hippocampus: p=0.008 control vs hexameric Aβ; in 5xFAD cortex: p=0.02 control vs hexameric Aβ).

### Insight into the cellular pathways and contribution of presenilins to the formation of cell-derived hexameric Aβ

Together, our results support that the identified cell-derived hexameric Aβ assembly has nucleating and seeding properties in the amyloid pathology. We aimed at understanding the cellular context in which this specific assembly is formed, and more precisely the contribution of PS1- and PS2-dependent γ-secretases to the formation of pathological Aβ assemblies. Previous studies demonstrated that PS1 and PS2 have differential substrate specificities [32, 37] and that several factors, including their specific subcellular localization [39], can orientate their amyloidogenic processing activity to the production of more or less aggregation-prone Aβ isoforms.

Hence, we investigated the formation of the assembly of interest in C99 expressing cells lacking PS1 or PS2. In order to use a homogenous system to evaluate processing of the human C99 substrate by human γ-secretases, we developed stable PS1 and PS2 *knockdown* (KD) neuron-derived cell lines (SH-SY5Y cells). SH-SY5Y readily produced the hexameric Aβ assembly in our conditions (Fig2C). Cells were transfected with a CRISPR-Cas9 expression system targeting either *PSEN1* or *PSEN2* genes, and selected using fluorescent (FACS) and antibiotic resistance double-selections. Scrambled (S) target sequences for both the *PSEN1* and *PSEN2* genes were used for the development of control cell lines. After sub-cloning, the expression of both PSs was verified by Western blotting (Fig6A) which showed a 44.7% and 63.2% reduction of PS1 and PS2 protein levels respectively. We first assessed the ability of the KD cells to perform the initial cleavage of the C99 substrate at the ε-site, releasing APP intracellular domain (AICD). The AICD release from a tagged C99-GVP substrate was measured by a Gal4 reporter gene assay, as described previously [37, 48]. Results revealed an efficient cleavage of the construct in both the PS1-KD and PS2-KD cells, when compared to PS1-S and PS2-S respectively, suggesting that neither of the *knockdown* performed here affect the ability to ensure substrate cleavage. We investigated the profile of Aβ production in these cell lines by Western blotting. Results indicated that the reduction in PS1 levels had no significant effect on the profile of Aβ produced inside or outside of the cell (Fig6B), with no significant decrease in monomeric Aβ_40_ or Aβ_42_ measured in culture media. This is quite an unexpected observation, that could be explained by the observation that only 50% of PS1 *knockdown* could be achieved in our model. The remaining PS1-dependent γ-secretase activity could be sufficient to efficiently process APP-derived substrates. However, while the formation of intracellular hexameric Aβ was similar between PS2-KD and corresponding control cells, detection of intracellular monomeric Aβ was lost when PS2 expression was reduced. Concomitantly, the extracellular Aβ assembly profile was altered in PS2-KD cells, with an increase in monomeric form (observed both by Western blotting and ECLIA) and an acute decrease in hexameric Aβ, suggesting that the extracellular release of hexameric Aβ is dependent on the presence of PS2 (Fig6C). This would illustrate that the absence of PS2 favors the accumulation of monomeric extracellular Aβ, but leads to decreased intracellular Aβ forms and extracellular aggregates. In other words, PS2-dependent γ-secretases could generate aggregation-prone intracellular Aβ, that is eventually released as an aggregate in the extracellular space. To note, PS2 [32, 39], as well as APP and intermediate fragments of its metabolism [58, 59], were previously found in endolysosomal compartments and extracellular vesicles (EVs). We examined whether hexameric Aβ assemblies found outside the cells were disposed of through EVs release, stemming from intracellular vesicular compartments. To discriminate whether Aβ species present in the media were compartmentalized in EVs, we performed a specific ultracentrifugation procedure to separate EVs from soluble proteins in the media of PS1-S, PS1-KD, PS2-S and PS2-KD cells. The efficiency of the separation was confirmed by the Europium-immunoassay with, in the EVs, significantly increased levels of inclusions markers CD9, CD63 and CD81 and lower content of the exclusion marker GM130 (Fig7A, left panel). The specific enrichment of inclusion markers due to higher content in proteins was ruled out since whole-protein assay showed larger protein amounts in soluble than EVs fractions (Fig7A, right panel). Finally, EVs size distribution was similar between all conditions but the number of EVs was higher in PS1-KD and PS2-KD as compared to PS1-S and PS2-S, although not significant except for PS2-S vs PS2-KD (Fig7B). Importantly, extracellular monomeric Aβ was found exclusively in the soluble proteins fraction while hexameric Aβ was confined exclusively in vesicles (Fig7C), as was very recently reported with Aβ oligomeric species [49]. C99 was decreased and hexameric Aβ assembly almost disappeared in PS2-KD EVs, indicating that PS2 plays a critical role in the extracellular release of this specific Aβ assembly that displays seeding properties.

**Fig6.**
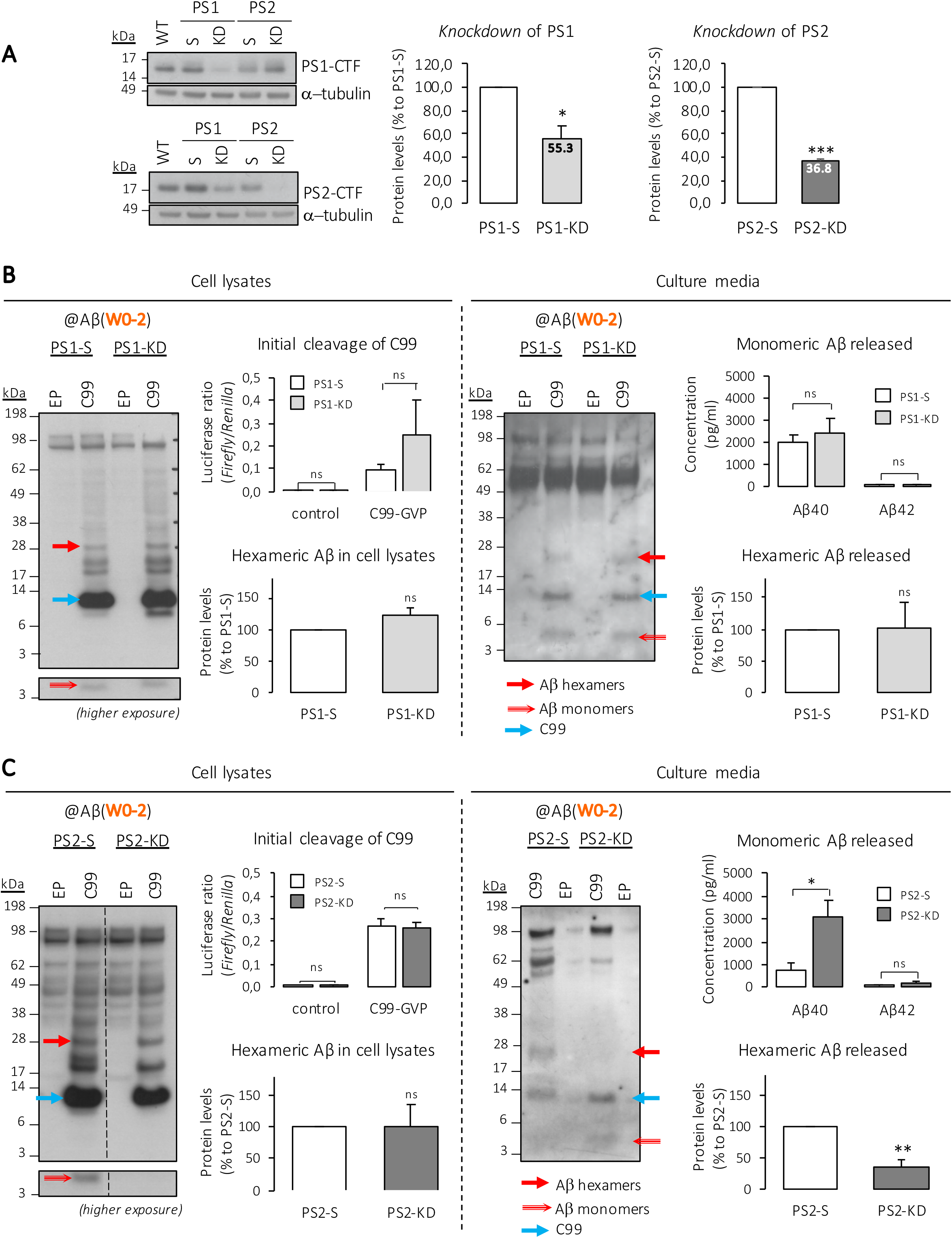
Contribution of presenilins to the production of hexameric Aβ. **A.** SH-SY5Y *knockdown* (KD) cell lines were generated using CRISPR-Cas9, with guide RNA vectors targeting either human *PSEN1* (PS1-KD) or *PSEN2* (PS2-KD) genes. Control cells were transfected with respective scrambled sequences. Left, a representative Western blot; middle and right, quantitative decrease in PS1 and PS2 protein levels in KD compared to S cells. N=3. One-sample *t* test with hypothetical value set as 100: *=p<0.05, ***=p<0.001 (S vs KD, in PS1: p=0.03; in PS2: p=0.0001). WT=wild-type, S=scramble. **B, C.** Initial cleavage ability was monitored by a reporter gene assay. The release of APP intracellular domain (AICD) from a tagged C99-GVP substrate was measured by the Gal4-*Firefly* reporter gene. Results are represented as *Firefly/Renilla* luciferases ratios, with *Renilla* serving as a transfection-efficiency control. The profile of Aβ production was assessed after transfection with either an empty plasmid (EP) or C99, using Western blotting and ECLIA immunoassay, in PS1-KD vs PS1-S (in **B.**) and in PS2-KD vs PS2-S (in **C.**). Dashed line indicates that proteins were run on the same gel, but lanes are not contiguous. Luciferase assays (initial cleavage of C99): N=4 each, one-way ANOVA with Tukey’s multiple comparison test: ns=non-significant (S vs KD, in PS1 control: p>0.99; in PS1 C99-GVP: p=0.10; in PS2 control: p=0.99; in PS2 C99-GVP: p=0.99). Western blots quantitative analyses (hexameric Aβ, in cell lysates and released, relative to C99 and % to S): N=3 each, one-sample *t* test with hypothetical value set as 1: ns=non-significant, **=p<0.01 (S vs KD, in PS1 cell lysates: p=0.16; in PS1 media: p=0.97; in PS2 cell lysates: p=0.99; in PS2 media: p=0.01). ECLIA assays (monomeric Aβ released): N=5 each, Mann-Whitney test: ns=non-significant, *=p<0.05 (S vs KD, in PS1 Aβ_40_: p>0.99; in PS1 Aβ_42_: p>0.99; in PS2 Aβ_40_: p=0.04; in PS2 Aβ_42_: p=0.12).

**Fig7.**
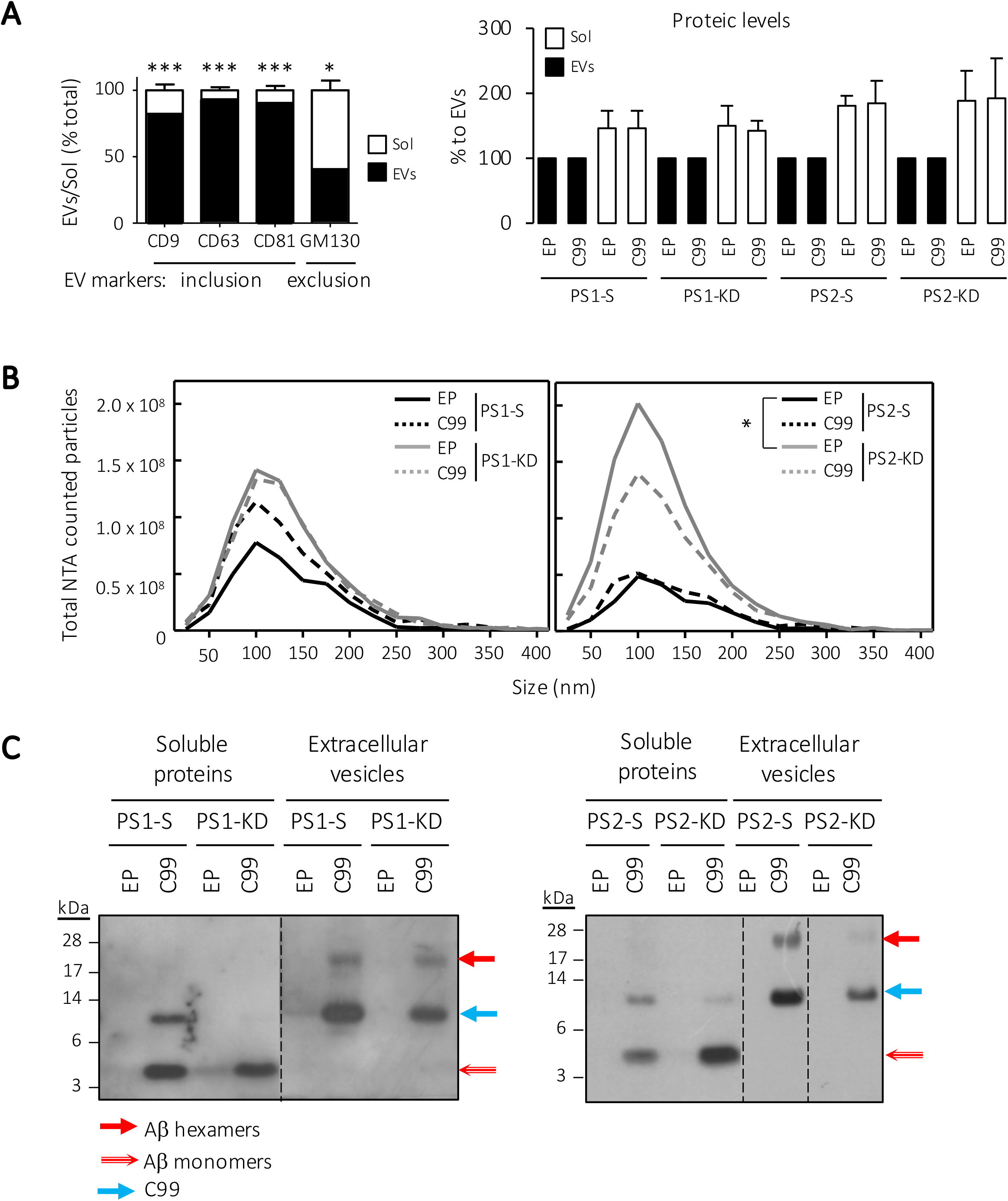
Localization of hexameric Aβ in extracellular vesicles (EVs). Media of PS1-S, PS1-KD, PS2-S and PS2-KD cells underwent an ultracentrifugation process to separate putative enrichment of EVs, in pellet, from soluble proteins. **A.** The efficiency of EV isolation was confirmed by plate-based Europium-immunoassay (left panel) showing an enrichment of EV inclusion markers CD9, CD63 and CD81 while EV exclusion marker GM130 was lower in EVs as compared to soluble fractions. N=6 independent experiments. Mann-Whitney test: *=p<0.05, ***=p<0.001 (CD9 (n=22), CD63 (n=22) and CD81 (n=18): p<0.0001; GM130 (n=19): p=0.0362). Quantification of protein levels by BCA (right panel) showed larger protein amounts in soluble than EVs fractions, ruling out the specific enrichment of inclusion markers due to higher content in proteins. **B.** EVs ultracentrifugation pellets were counted for number of particles of 70-400nm by nanoparticle tracking analysis (NTA). n=9 in N=3 independent experiments. Two-way ANOVA with Bonferroni’s multiple comparison: *=p<0.05 (PS2-KD EP vs PS2-S EP: p<0.05). **C.** Both EVs and soluble extracts were monitored by Western blotting with the W0-2 antibody. Dashed lines indicate that proteins were run on the same gel, but lanes are not contiguous. EP=empty plasmid, EV(s)=extracellular vesicle(s), Sol=soluble proteins fraction.

## Discussion

The identification of fibrillar Aβ as the main component of amyloid plaques present in AD has led to the extensive investigation of the Aβ self-assembly process and, as a result, to the identification of many intermediate oligomeric assemblies of Aβ. Today, it is widely agreed that such non-fibrillar soluble assemblies exert a widespread neurotoxic effect and should be the main target of therapeutic prevention or intervention. Among Aβ oligomers, hexameric Aβ has repeatedly been identified as a key assembly *in vitro* [15, 22, 60, 61]. Our study shows the identification of a specific ^~^28kDa assembly in a wide range of models. Using CHO cells, we were able to confirm the nature of this assembly as hexameric Aβ_42_. As mentioned previously, many studies, most of which were performed on synthetic preparations of Aβ, have shed light on the importance of hexameric Aβ as a crucial nucleating step in the process of Aβ self-assembly [15, 23, 24, 62]. However, such studies placing hexameric Aβ at the core of amyloid pathology have, to the best of our knowledge, not been able to characterize the production of Aβ hexamers in a cellular environment, or to assess its toxic potential and relevance to AD pathogenesis.

Here, we report for the first time the identification of hexameric Aβ assemblies in brain extracts from a well-characterized amyloid mouse model (5xFAD) as well as in the CSF of AD patients. The question of whether this assembly is strictly speaking formed by six Aβ_42_ monomers associated together would require extensive analytical biochemistry investigation, which are of real interest, but beyond the scope of this study. Nevertheless, the Aβ assembly we identified and purified by GELFrEE electrophoresis (i) has an apparent molecular weight around 28kDa, (ii) is recognized by W0-2, 6E10 and anti-Aβ_42_ antibodies but not by an APP-C-ter antibody and (iii) decreases upon PS2 *knockdown*. Together, this clearly indicates that the detected Aβ assembly contains Aβ_42_ and has the expected properties of a hexamer.

Our observations conclusively place the assembly of interest at the center of amyloid pathogenesis. In the 5xFAD model, the presence of hexameric Aβ followed a regional pattern of progression that corresponds to the neuropathological staging of human AD pathogenesis described by Braak and Braak [63]. Indeed, the ^~^28kDa assembly was detected first in the hippocampus, as early as 2-month-old, and further spread to the cortex starting from 3 to 6-month-old. On the contrary, amyloid plaques formation in the 5xFAD mice have previously been reported to appear first in deep layers of the cortex and in the subiculum, and to later spread to the hippocampus as mice aged [28]. As the spreading properties of hexameric Aβ are likely to depend on the initial site where they are formed, it would be of interest to investigate the aggravation of amyloid pathology upon injection of cell-derived hexameric Aβ in the cortical areas. The pattern of hexameric Aβ detection reported here is also of particular importance considering the urgent need to identify early biomarkers in the development of A β pathology. In line with this, the detection of the ^~^28kDa assembly in CSF extracted from human patients diagnosed with pre-clinical AD and symptomatic AD supports its implication in the onset and development of the pathology. More importantly, its presence as a soluble entity in CSF is a strong argument to consider it as a novel detectable biomarker.

Furthermore, the isolation of hexameric Aβ from our CHO cell model has also allowed the characterization of its detrimental properties. The treatment of primary WT neurons with 1μM of cell-derived hexameric Aβ for 24h did not indicate any cytotoxic effect. These experimental conditions mimic a cellular environment comparable to the early phase of oligomeric Aβ pathology, as reported in several studies [64–67]. Still, as concentrations of oligomeric Aβ_42_ have been reported to reach concentrations of up to 3μM inside AD-affected neurons [68], we also tested a 5μM concentration in our assay. Yet, we did not observe any increase in cell death when comparing neurons treated with cell-derived hexameric Aβ to control-treated neurons. Interestingly, another study from our group based on ThT assays and Western blotting revealed a very stable behavior of the hexameric assembly when incubated by itself *in vitro*, unable to further aggregate [29]. The lack of direct harmful effects on neurons is in line with the fact that (i) the process of Aβ self-assembly is thought to be vital in mediating cytotoxicity [69, 70], and that (ii) the pathological properties of Aβ oligomers could rely not only on their synaptotoxic effects but also on their seeding properties, propagating amyloid pathology throughout the brain parenchyma.

In line with the observation that cell-derived hexameric Aβ does not appear cytotoxic by itself, and considering its suggested role as a nucleus for Aβ self-assembly, we studied whether the harmful potential of Aβ hexamers could be unraveled when pre-existing Aβ species are present. Cytotoxic assay on primary neurons derived from transgenic 5xFAD mice was performed. It was previously reported that cultured neurons from AD transgenic animal models can reflect AD phenotypes *in vitro* [55–57]. We observed a significant increase in the proportion of cell death when neurons were treated with 5 μM of cell-derived hexameric Aβ. This suggests that this specific assembly can indeed exert a toxic effect in a FAD context, when Aβ to seed is likely present. To further assess this hypothesis, we performed stereotaxic injections of isolated cell-derived hexameric Aβ in 5xFAD mouse brains and followed Aβ deposition, in parallel to WT injected mice. The results obtained in WT mice suggest that the identified assembly is not able to induce Aβ deposition *in vivo* by itself. This is in agreement with the absence of cytotoxicity on WT primary neurons reported above. To note, Aβ deposition *in vivo* was assessed in a time-frame of 30 days. One cannot exclude that pathogenic mechanisms might take place upon longer incubation time. In addition, the absence of cytotoxicity or Aβ deposits formation by cell-derived hexameric Aβ *per se* does not exclude its ability to cause changes in primary neurons or in the brain function apart from toxicity or amyloid deposition. Indeed, cellular dysfunctions or alterations of neuronal connectivity might be present in our WT models and simply not yet sufficient to cause cytotoxicity or amyloidosis. Additionally, the absence of Aβ deposits after injection of cell-derived hexameric Aβ in the brain of WT mice also does not exclude the possibility that the stable seeds injected might not directly cause amyloidosis in the injected animals, but persist in the brain and retain pathogenic activity, as was previously shown with second-transmission studies [5].

A major observation in this study is the ability of cell-derived hexameric Aβ to act as a seeding nucleus and cause both cytotoxicity in primary neurons and aggravation of Aβ deposition in the brain when using transgenic 5xFAD mice. These mice express five familial AD mutations that together trigger Aβ_42_ overproduction and result in a rapid and severe development of amyloid pathology [28, 50]. 5xFAD mice therefore represented a useful model to assess the nucleating hypothesis *in vivo* in a reasonable timeframe. An earlier onset of Aβ aggregation in this model was previously reported upon single intracerebral injection of brain homogenates containing oligomeric A β, following a prion-like seeding mechanism [71]. Aβ oligomers have been further suggested as early initiating actors of the seeding process, as their depletion by passive immunization delays Aβ aggregation and lead to a transient reduction of seed-induced Aβ deposition [72]. Here, our study has the advantage of focusing on a specific cell-derived Aβ aggregate, that not only is an oligomeric entity of its own but also suspected, based on *in vitro* studies, to be heavily involved in the processes of nucleation and seeding [15, 29]. We chose to perform intracerebral injections at 2 months of age, when amyloid deposition begins in the 5xFAD mice [28]. The significant increase of Aβ deposition observed in hexamer-injected hemispheres suggests that hexameric Aβ is indeed able to promote amyloidosis. As very recent *in vitro* studies from our group revealed the ability of the isolated Aβ hexamers to drive the aggregation of synthetic monomers of Aβ [29], it is likely that the enhancement in Aβ aggregation observed here relies on the same process of nucleation, with Aβ hexamers serving as a template for aggregation. The greater increase of Aβ deposition in the hippocampus when compared to the cortex of the injected mice supports this hypothesis, as hexameric Aβ is likely to seed and promote Aβ aggregation to a higher rate at the site of injection. Still, the significant aggravation of deposition in the cortex also suggests that hexameric Aβ is able to spread throughout the brain, as rapidly as in 30 days. Together, these observations strongly suggest that hexameric Aβ accounts for a key factor in the self-assembly process, which is able to promote amyloidosis and might serve, when naturally present, as an early biomarker for Aβ deposition.

Following this important hypothesis, we sought to understand the cellular context in which hexameric Aβ is produced, in hope of unraveling potential therapeutic targets. In particular, we assessed the respective involvement of both presenilins in its production. Indeed, the distinct subcellular localizations [39] and differential substrate specificities [37] of the two types of PSs regulate the production of different Aβ pools. Production of Aβ can differ considerably between cellular compartments [39]. Pathological Aβ formation is related to dysfunction of the endocytic pathway, and PS1 and PS2 are differentially distributed between the secretory compartments and the late endosomes/lysosomes to which PS2 is shuttled [39]. Our results showed the absence of any significant change in the processing of C99 or the release of Aβ when cells have a nearly 50% reduction of PS1 protein levels. This was rather surprising regarding the previous reports on the preponderant importance of PS1 for γ-secretase substrates cleavage and overall Aβ production [33–37]. One can imagine, as mentioned above, that the reduction of PS1 protein levels is not significant enough to observe any effect. PS1 *knockdown* might induce compensatory mechanisms and still ensure its primary function even when its protein levels are reduced by half. In any case, results reported here bring interesting information regarding the production of hexameric Aβ at the cellular level. Indeed, in the PS2 *knockdown* cells, the reduction of PS2 protein levels by just over 60% was sufficient to cause clear changes in Aβ production. While the initial cleavage of the C99 construct (ε-site) remained unaffected, the production of monomeric Aβ was strongly diminished inside the cell and, in an opposite manner, strongly increased in the medium. In particular, the Aβ_40_ isoform was increased in the extracellular medium of PS2-KD cells. Hexameric Aβ levels were unchanged in the cell lysates but strongly diminished in the extracellular environment. Importantly, we could discriminate for the first time that the extracellular hexameric A β was almost exclusively enriched in EVs while monomeric Aβ was identified solely in the soluble fraction. Notice that, although the amount of EVs tended to increase upon *knockdown* of PS1/PS2, it did not seem to impact Aβ fate. As PS1 and PS2 have been respectively shown to produce the extra- and intracellular pools of Aβ [39], the direct release of soluble Aβ outside the cellular environment is likely to rely mostly on the action of PS1, while the intracellular pool is likely to be mainly dependent of PS2 activity in vesicular compartments. Our observations suggest that a reduction of PS2 levels strongly influences the production of Aβ from the C99 fragment. Monomeric Aβ_42_ is likely to completely aggregate into hexamers, while monomeric Aβ_40_ might not able to aggregate, causing the observed increase in the detection of this isoform outside the cell. Importantly, this is supported by observations reported in a biochemical *in vitro* study [29], showing the isolation of monomeric Aβ from the medium of C99-expressing CHO cells and identifying it as Aβ_40_, not forming any aggregates when followed over 48h.

Together, our results in both *knockdown* cell lines drive towards the emergence of a new hypothesis according to which extracellular hexameric Aβ might be exclusively released in EVs emerging from the intracellular pool of Aβ and likely produced through the activity of PS2. The identification of such a specific role for PS2 in the release of hexameric Aβ that might exert intercellular nucleating effects is of particular importance. To note, PS1 and PS2 were reported to exert different sensibilities to γ-secretase inhibitors [37]. This is quite promising in the hope of re-evaluating Aβ modulators and developing therapeutic agents targeting a specific γ-secretase activity depending on the PS present in the complex. Importantly, the intracellular pool of Aβ, generated by PS2, has been repeatedly associated with the progression of AD [73–76]. FAD mutations in *PSEN2* have been shown to dramatically increase the proportion of longer length Aβ intracellularly, accelerating its assembly. Further, a subset of familial mutations on *PSEN1* have been reported to shift the localization of the PS1 protein to fit that of PS2 [39]. Thus, it is likely that the production of aggregation-prone Aβ inside intracellular compartments and its resulting accumulation and excretion are enhanced in the context of AD.

## Conclusions

Altogether, our findings have shed light on a particular cell-derived Aβ assembly that likely corresponds to an Aβ_42_ hexamer. Combining *in vitro* and *in vivo* approaches, we have revealed an absence of detrimental effects of cell-derived hexameric Aβ by itself, but its capacity to induce cytotoxicity and aggravate amyloid deposition when there is Aβ to seed at disposal. An insight in cellular mechanisms at stake suggests a stronger correlation of PS2 with the formation of this particular Aβ oligomer, in line with previous reports linking the restricted location of PS2 in acidic compartments to the production of more aggregation-prone Aβ.

## Declarations

### Ethics approval and consent to participate

All animal experiments were performed with the approval of the UCLouvain Ethical Committee for Animal Welfare (reference 2018/UCL/MD/011). Human cerebrospinal fluids were collected as part of clinical analyses performed at Cliniques Universitaires Saint-Luc (UCL, Brussels, Belgium). Symptomatic non-AD patients signed an internal regulatory document, stating that residual samples used for diagnostic procedures can be used for retrospective academic studies, without any additional informed consent (ethics committee approval: 2007/10SEP/233). AD patients participated to a specific study referenced UCL-2016-121 (Eudra-CT: 2018-003473-94).

### Consent for publication

All authors have given consent for publication.

### Availability of data and materials

All datasets generated and analyzed during this study are included in this published article and its supplementary information files. Materials are available upon request.

### Competing interests

The authors declare that the research was conducted in the absence of any commercial or financial relationships that could be construed as a potential conflict of interest.

### Funding

This work was supported by a grant of the Belgian F.N.R.S FRIA (Fonds National pour la Recherche Scientifique) and a grant of UCLouvain Fonds du Patrimoine to CV. Funding to PKC is acknowledged from SAO-FRA Alzheimer Research Foundation, Fondation Louvain and Queen Elisabeth Medical Research Foundation (FMRE to PKC and LQ). The work was supported by funds from FNRS grant PDRT.0177.18 to PKC and LQ.

### Authors’ contributions

CV designed and performed experiments, analyzed and interpreted data, and wrote the manuscript. DMV performed experiments and analyzed data. SC and NS provided neuronal cultures and analyzed data. LD’A performed nanoparticle tracking and Europium-immunoassay analyses on isolated EVs, and helped with the analysis of related data. FP participated in experiment design and analysis. VVP and BH together provided the human CSF specimens. BH provided significant input in the interpretation of clinical data. LQ provided the GELFrEE technique and contributed to data collection. PKC designed and supervised the research project and contributed to interpretation of data. All authors revised and approved the final manuscript.

## Abbreviations

Aβ: β-amyloid
AD: Alzheimer’s disease
AICD: APP intracellular domain
APP: amyloid precursor protein
CHO: Chinese hamster ovary
CRISPR: clustered regularly interspaced short palindromic repeats
Cryo-EM: cryo-electron microscopy
CSF: cerebrospinal fluid
CTF: C-terminal fragment
HEK293: human embryonic kidney
KD: *knockdown*
MCI: mild cognitive impairment
PET: positron-emission tomography
*PSEN*: presenilin (gene)
PS: presenilin (protein)
S: scrambled
ThT: Thioflavin T
WT: wild-type.

## Acknowledgments

We thank Esther Paître, Sarah Houben and Pierre Burguet for technical support. We are grateful to Jean-Noël Octave, Nathalie Pierrot and Jean-Pierre Brion for providing scientific input as well as materials and animal models.

## Additional files

**Additional File 1.**

Format: .pdf

Title: **Detection of hexameric Aβ_42_ by the anti-Aβ clone 6E10 primary antibody.**

Description: Cell lysates and media of CHO cells were processed as in Fig1. Detection with the 6E10 clone targeting human Aβ revealed the same ^~^28kDa bands as W0-2 when cells are expressing either C42 or C99, reinforcing their identification as Aβ assemblies. Dashed lines indicate that proteins were run on the same gel, but lanes are not contiguous.

**Additional File 2.**

Format: .xls

Title: **Clinical analyses performed on the patients used in this study.**

Description: Non-AD (1258 and 1506), pre-clinical AD (1523, 1268 and 1556), and symptomatic AD (1272, 1329 and 1633) patients were monitored for memory impairments by the Mini-Mental State Examination (MMSE). Their CSF was collected for measurement of the two biomarkers of AD; Aβ and Tau. PET imaging using amyloid and/or Tau specific ligands was conducted in patients where the measurements exceeded pathological threshold. +=positive, N/A=non acquired.

## Notes

### Competing Interest Statement

The authors have declared no competing interest.

## References

1. Haass C, Kaether C, Thinakaran G, Sisodia S: Trafficking and proteolytic processing of APP. Cold Spring Harb Perspect Med 2012, 2:a006270.

2. Takami M, Nagashima Y, Sano Y, Ishihara S, Morishima-Kawashima M, Funamoto S, Ihara Y: gamma-Secretase: successive tripeptide and tetrapeptide release from the transmembrane domain of beta-carboxyl terminal fragment. J Neurosci 2009, 29:13042–13052.

3. Benilova I, Karran E, De Strooper B: The toxic Aβ oligomer and Alzheimer’s disease: an emperor in need of clothes. Nat Neurosci 2012, 15:349–357.

4. Almeida ZL, Brito RMM: Structure and Aggregation Mechanisms in Amyloids. Molecules 2020, 25.

5. Ye L, Fritschi SK, Schelle J, Obermüller U, Degenhardt K, Kaeser SA, Eisele YS, Walker LC, Baumann F, Staufenbiel M, Jucker M: Persistence of Aβ seeds in APP null mouse brain. Nat Neurosci 2015, 18:1559–1561.

6. Sengupta U, Nilson AN, Kayed R: The Role of Amyloid-β Oligomers in Toxicity, Propagation, and Immunotherapy. EBioMedicine 2016, 6:42–49.

7. Ferrone F: Analysis of protein aggregation kinetics. Methods Enzymol 1999, 309:256–274.

8. Serpell LC: Alzheimer’s amyloid fibrils: structure and assembly. Biochim Biophys Acta 2000, 1502:16–30.

9. Fu Z, Aucoin D, Davis J, Van Nostrand WE, Smith SO: Mechanism of Nucleated Conformational Conversion of Aβ42. Biochemistry 2015, 54:4197–4207.

10. Lambert MP, Barlow AK, Chromy BA, Edwards C, Freed R, Liosatos M, Morgan TE, Rozovsky I, Trommer B, Viola KL, et al: Diffusible, nonfibrillar ligands derived from Abeta1-42 are potent central nervous system neurotoxins. Proc Natl Acad Sci U S A 1998, 95:6448–6453.

11. Sunde M, Blake CC: From the globular to the fibrous state: protein structure and structural conversion in amyloid formation. Q Rev Biophys 1998, 31:1–39.

12. Klein WL, Krafft GA, Finch CE: Targeting small Abeta oligomers: the solution to an Alzheimer’s disease conundrum? Trends Neurosci 2001, 24:219–224.

13. Chen GF, Xu TH, Yan Y, Zhou YR, Jiang Y, Melcher K, Xu HE: Amyloid beta: structure, biology and structure-based therapeutic development. Acta Pharmacol Sin 2017, 38:1205–1235.

14. Cline EN, Bicca MA, Viola KL, Klein WL: The Amyloid-β Oligomer Hypothesis: Beginning of the Third Decade. J Alzheimers Dis 2018, 64:S567–s610.

15. Roychaudhuri R, Yang M, Hoshi MM, Teplow DB: Amyloid beta-protein assembly and Alzheimer disease. J Biol Chem 2009, 284:4749–4753.

16. Walsh DM, Klyubin I, Fadeeva JV, Rowan MJ, Selkoe DJ: Amyloid-beta oligomers: their production, toxicity and therapeutic inhibition. Biochem Soc Trans 2002, 30:552–557.

17. Lesné S, Koh MT, Kotilinek L, Kayed R, Glabe CG, Yang A, Gallagher M, Ashe KH: A specific amyloid-beta protein assembly in the brain impairs memory. Nature 2006, 440:352–357.

18. Townsend M, Shankar GM, Mehta T, Walsh DM, Selkoe DJ: Effects of secreted oligomers of amyloid beta-protein on hippocampal synaptic plasticity: a potent role for trimers. J Physiol 2006, 572:477–492.

19. Shankar GM, Li S, Mehta TH, Garcia-Munoz A, Shepardson NE, Smith I, Brett FM, Farrell MA, Rowan MJ, Lemere CA, et al: Amyloid-beta protein dimers isolated directly from Alzheimer’s brains impair synaptic plasticity and memory. Nat Med 2008, 14:837–842.

20. Lesné SE, Sherman MA, Grant M, Kuskowski M, Schneider JA, Bennett DA, Ashe KH: Brain amyloid-β oligomers in ageing and Alzheimer’s disease. Brain 2013, 136:1383–1398.

21. Müller-Schiffmann A, Herring A, Abdel-Hafiz L, Chepkova AN, Schäble S, Wedel D, Horn AH, Sticht H, de Souza Silva MA, Gottmann K, et al: Amyloid-β dimers in the absence of plaque pathology impair learning and synaptic plasticity. Brain 2016, 139:509–525.

22. Wolff M, Zhang-Haagen B, Decker C, Barz B, Schneider M, Biehl R, Radulescu A, Strodel B, Willbold D, Nagel-Steger L: Aβ42 pentamers/hexamers are the smallest detectable oligomers in solution. Sci Rep 2017, 7:2493.

23. Cernescu M, Stark T, Kalden E, Kurz C, Leuner K, Deller T, Göbel M, Eckert GP, Brutschy B: Laser-induced liquid bead ion desorption mass spectrometry: an approach to precisely monitor the oligomerization of the β-amyloid peptide. Anal Chem 2012, 84:5276–5284.

24. Österlund N, Moons R, Ilag LL, Sobott F, Gräslund A: Native Ion Mobility-Mass Spectrometry Reveals the Formation of β-Barrel Shaped Amyloid-β Hexamers in a Membrane-Mimicking Environment. J Am Chem Soc 2019, 141:10440–10450.

25. Deshpande A, Mina E, Glabe C, Busciglio J: Different conformations of amyloid beta induce neurotoxicity by distinct mechanisms in human cortical neurons. J Neurosci 2006, 26:6011–6018.

26. Gellermann GP, Byrnes H, Striebinger A, Ullrich K, Mueller R, Hillen H, Barghorn S: Abeta-globulomers are formed independently of the fibril pathway. Neurobiol Dis 2008, 30:212–220.

27. Decock M, Stanga S, Octave JN, Dewachter I, Smith SO, Constantinescu SN, Kienlen-Campard P: Glycines from the APP GXXXG/GXXXA Transmembrane Motifs Promote Formation of Pathogenic Aβ Oligomers in Cells. Front Aging Neurosci 2016, 8:107.

28. Oakley H, Cole SL, Logan S, Maus E, Shao P, Craft J, Guillozet-Bongaarts A, Ohno M, Disterhoft J, Van Eldik L, et al: Intraneuronal beta-amyloid aggregates, neurodegeneration, and neuron loss in transgenic mice with five familial Alzheimer’s disease mutations: potential factors in amyloid plaque formation. J Neurosci 2006, 26:10129–10140.

29. Vadukul DM, Vrancx C, Burguet P, Contino S, Suelves N, Serpell LC, Quinton L, Kienlen-Campard P: Cell-derived hexameric β-amyloid: a novel insight into composition, self-assembly and nucleating properties. Submitted bioRxiv doi: 101101/20201215422916 2020.

30. De Strooper B: Aph-1, Pen-2, and Nicastrin with Presenilin generate an active gamma-Secretase complex. Neuron 2003, 38:9–12.

31. Sato T, Diehl TS, Narayanan S, Funamoto S, Ihara Y, De Strooper B, Steiner H, Haass C, Wolfe MS: Active gamma-secretase complexes contain only one of each component. J Biol Chem 2007, 282:33985–33993.

32. Dehury B, Tang N, Blundell TL, Kepp KP: Structure and dynamics of γ-secretase with presenilin 2 compared to presenilin 1. RSC Advances 2019, 9:20901–20916.

33. De Strooper B, Saftig P, Craessaerts K, Vanderstichele H, Guhde G, Annaert W, Von Figura K, Van Leuven F: Deficiency of presenilin-1 inhibits the normal cleavage of amyloid precursor protein. Nature 1998, 391:387–390.

34. Frånberg J, Svensson AI, Winblad B, Karlström H, Frykman S: Minor contribution of presenilin 2 for γ-secretase activity in mouse embryonic fibroblasts and adult mouse brain. Biochem Biophys Res Commun 2011, 404:564–568.

35. Pintchovski SA, Schenk DB, Basi GS: Evidence that enzyme processivity mediates differential Aβ production by PS1 and PS2. Curr Alzheimer Res 2013, 10:4–10.

36. Xia D, Watanabe H, Wu B, Lee SH, Li Y, Tsvetkov E, Bolshakov VY, Shen J, Kelleher RJ, 3rd: Presenilin-1 knockin mice reveal loss-of-function mechanism for familial Alzheimer’s disease. Neuron 2015, 85:967–981.

37. Stanga S, Vrancx C, Tasiaux B, Marinangeli C, Karlström H, Kienlen-Campard P: Specificity of presenilin-1-and presenilin-2-dependent γ-secretases towards substrate processing. J Cell Mol Med 2018, 22:823–833.

38. Meckler X, Checler F: Presenilin 1 and Presenilin 2 Target γ-Secretase Complexes to Distinct Cellular Compartments. J Biol Chem 2016, 291:12821–12837.

39. Sannerud R, Esselens C, Ejsmont P, Mattera R, Rochin L, Tharkeshwar AK, De Baets G, De Wever V, Habets R, Baert V, et al: Restricted Location of PSEN2/γ-Secretase Determines Substrate Specificity and Generates an Intracellular Aβ Pool. Cell 2016, 166:193–208.

40. Lee S, Mankhong S, Kang JH: Extracellular Vesicle as a Source of Alzheimer’s Biomarkers: Opportunities and Challenges. Int J Mol Sci 2019, 20.

41. Wiedenheft B, Sternberg SH, Doudna JA: RNA-guided genetic silencing systems in bacteria and archaea. Nature 2012, 482:331–338.

42. Hsu PD, Lander ES, Zhang F: Development and applications of CRISPR-Cas9 for genome engineering. Cell 2014, 157:1262–1278.

43. Kienlen-Campard P, Tasiaux B, Van Hees J, Li M, Huysseune S, Sato T, Fei JZ, Aimoto S, Courtoy PJ, Smith SO, et al: Amyloidogenic processing but not amyloid precursor protein (APP) intracellular C-terminal domain production requires a precisely oriented APP dimer assembled by transmembrane GXXXG motifs. J Biol Chem 2008, 283:7733–7744.

44. Huysseune S, Kienlen-Campard P, Hébert S, Tasiaux B, Leroy K, Devuyst O, Brion JP, De Strooper B, Octave JN: Epigenetic control of aquaporin 1 expression by the amyloid precursor protein. Faseb j 2009, 23:4158–4167.

45. Teunissen CE, Petzold A, Bennett JL, Berven FS, Brundin L, Comabella M, Franciotta D, Frederiksen JL, Fleming JO, Furlan R, et al: A consensus protocol for the standardization of cerebrospinal fluid collection and biobanking. Neurology 2009, 73:1914–1922.

46. Hage S, Stanga S, Marinangeli C, Octave JN, Dewachter I, Quetin-Leclercq J, Kienlen-Campard P: Characterization of Pterocarpus erinaceus kino extract and its gamma-secretase inhibitory properties. J Ethnopharmacol 2015, 163:192–202.

47. Opsomer R, Contino S, Perrin F, Gualdani R, Tasiaux B, Doyen P, Vergouts M, Vrancx C, Doshina A, Pierrot N, et al: Amyloid Precursor Protein (APP) Controls the Expression of the Transcriptional Activator Neuronal PAS Domain Protein 4 (NPAS4) and Synaptic GABA Release. eNeuro 2020, 7.

48. Karlström H, Bergman A, Lendahl U, Näslund J, Lundkvist J: A sensitive and quantitative assay for measuring cleavage of presenilin substrates. J Biol Chem 2002, 277:6763–6766.

49. Perrin F, Papadopoulos N, Suelves N, Opsomer R, Vadukul DM, Vrancx C, Smith SO, Vertommen D, Kienlen-Campard P, Constantinescu SN: Dimeric Transmembrane Orientations of APP/C99 Regulate γ-Secretase Processing Line Impacting Signaling and Oligomerization. ISCIENCE 2020.

50. Eimer WA, Vassar R: Neuron loss in the 5XFAD mouse model of Alzheimer’s disease correlates with intraneuronal Aβ42 accumulation and Caspase-3 activation. Mol Neurodegener 2013, 8:2.

51. Doecke JD, Pérez-Grijalba V, Fandos N, Fowler C, Villemagne VL, Masters CL, Pesini P, Sarasa M: Total Aβ(42)/Aβ(40) ratio in plasma predicts amyloid-PET status, independent of clinical AD diagnosis. Neurology 2020, 94:e1580–e1591.

52. Gravina SA, Ho L, Eckman CB, Long KE, Otvos L, Jr., Younkin LH, Suzuki N, Younkin SG: Amyloid beta protein (A beta) in Alzheimer’s disease brain. Biochemical and immunocytochemical analysis with antibodies specific for forms ending at A beta 40 or A beta 42(43). J Biol Chem 1995, 270:7013–7016.

53. Jarrett JT, Berger EP, Lansbury PT, Jr.: The carboxy terminus of the beta amyloid protein is critical for the seeding of amyloid formation: implications for the pathogenesis of Alzheimer’s disease. Biochemistry 1993, 32:4693–4697.

54. Li X, Buxbaum JN: Transthyretin and the brain re-visited: is neuronal synthesis of transthyretin protective in Alzheimer’s disease? Mol Neurodegener 2011, 6:79.

55. Kim H, Kim B, Kim HS, Cho JY: Nicotinamide attenuates the decrease in dendritic spine density in hippocampal primary neurons from 5xFAD mice, an Alzheimer’s disease animal model. Mol Brain 2020, 13:17.

56. Mariani MM, Malm T, Lamb R, Jay TR, Neilson L, Casali B, Medarametla L, Landreth GE: Neuronally-directed effects of RXR activation in a mouse model of Alzheimer’s disease. Sci Rep 2017, 7:42270.

57. Noh H, Park C, Park S, Lee YS, Cho SY, Seo H: Prediction of miRNA-mRNA associations in Alzheimer’s disease mice using network topology. BMC Genomics 2014, 15:644.

58. Lauritzen I, Pardossi-Piquard R, Bourgeois A, Pagnotta S, Biferi MG, Barkats M, Lacor P, Klein W, Bauer C, Checler F: Intraneuronal aggregation of the β-CTF fragment of APP (C99) induces Aβ-independent lysosomal-autophagic pathology. Acta Neuropathol 2016, 132:257–276.

59. Evrard C, Kienlen-Campard P, Coevoet M, Opsomer R, Tasiaux B, Melnyk P, Octave JN, Buée L, Sergeant N, Vingtdeux V: Contribution of the Endosomal-Lysosomal and Proteasomal Systems in Amyloid-β Precursor Protein Derived Fragments Processing. Front Cell Neurosci 2018, 12:435.

60. Bitan G, Kirkitadze MD, Lomakin A, Vollers SS, Benedek GB, Teplow DB: Amyloid beta-protein (Abeta) assembly: Abeta 40 and Abeta 42 oligomerize through distinct pathways. Proc Natl Acad Sci U S A 2003, 100:330–335.

61. Roychaudhuri R, Yang M, Deshpande A, Cole GM, Frautschy S, Lomakin A, Benedek GB, Teplow DB: C-terminal turn stability determines assembly differences between Aβ40 and Aβ42. J Mol Biol 2013, 425:292–308.

62. Lendel C, Bjerring M, Dubnovitsky A, Kelly RT, Filippov A, Antzutkin ON, Nielsen NC, Härd T: A hexameric peptide barrel as building block of amyloid-β protofibrils. Angew Chem Int Ed Engl 2014, 53:12756–12760.

63. Braak H, Braak E: Neuropathological stageing of Alzheimer-related changes. Acta Neuropathol 1991, 82:239–259.

64. Nath S, Agholme L, Kurudenkandy FR, Granseth B, Marcusson J, Hallbeck M: Spreading of neurodegenerative pathology via neuron-to-neuron transmission of β-amyloid. J Neurosci 2012, 32:8767–8777.

65. Hallbeck M, Nath S, Marcusson J: Neuron-to-neuron transmission of neurodegenerative pathology. Neuroscientist 2013, 19:560–566.

66. Domert J, Rao SB, Agholme L, Brorsson AC, Marcusson J, Hallbeck M, Nath S: Spreading of amyloid-β peptides via neuritic cell-to-cell transfer is dependent on insufficient cellular clearance. Neurobiol Dis 2014, 65:82–92.

67. Sackmann C, Hallbeck M: Oligomeric amyloid-β induces early and widespread changes to the proteome in human iPSC-derived neurons. Sci Rep 2020, 10:6538.

68. Hashimoto M, Bogdanovic N, Volkmann I, Aoki M, Winblad B, Tjernberg LO: Analysis of microdissected human neurons by a sensitive ELISA reveals a correlation between elevated intracellular concentrations of Abeta42 and Alzheimer’s disease neuropathology. Acta Neuropathol 2010, 119:543–554.

69. Marshall KE, Vadukul DM, Dahal L, Theisen A, Fowler MW, Al-Hilaly Y, Ford L, Kemenes G, Day IJ, Staras K, Serpell LC: A critical role for the self-assembly of Amyloid-β1-42 in neurodegeneration. Sci Rep 2016, 6:30182.

70. Castelletto V, Ryumin P, Cramer R, Hamley IW, Taylor M, Allsop D, Reza M, Ruokolainen J, Arnold T, Hermida-Merino D, et al: Self-Assembly and Anti-Amyloid Cytotoxicity Activity of Amyloid beta Peptide Derivatives. Sci Rep 2017, 7:43637.

71. Morales R, Callegari K, Soto C: Prion-like features of misfolded Aβ and tau aggregates. Virus Res 2015, 207:106–112.

72. Katzmarski N, Ziegler-Waldkirch S, Scheffler N, Witt C, Abou-Ajram C, Nuscher B, Prinz M, Haass C, Meyer-Luehmann M: Aβ oligomers trigger and accelerate Aβ seeding. Brain Pathol 2020, 30:36–45.

73. Bayer TA, Wirths O: Intraneuronal Aβ as a trigger for neuron loss: ca n this be translated into human pathology? Biochem Soc Trans 2011, 39:857–861.

74. Friedrich RP, Tepper K, Rönicke R, Soom M, Westermann M, Reymann K, Kaether C, Fändrich M: Mechanism of amyloid plaque formation suggests an intracellular basis of Abeta pathogenicity. Proc Natl Acad Sci U S A 2010, 107:1942–1947.

75. Gouras GK, Tampellini D, Takahashi RH, Capetillo-Zarate E: Intraneuronal beta-amyloid accumulation and synapse pathology in Alzheimer’s disease. Acta Neuropathol 2010, 119:523–541.

76. Pensalfini A, Albay R, 3rd, Rasool S, Wu JW, Hatami A, Arai H, Margol L, Milton S, Poon WW, Corrada MM, et al: Intracellular amyloid and the neuronal origin of Alzheimer neuritic plaques. Neurobiol Dis 2014, 71:53–61.

