## Supplementary figures and images for "Mechanism of cellular production and *in vivo* seeding effects of hexameric β-amyloid assemblies"

### Additional File 1

#### Additional File 1

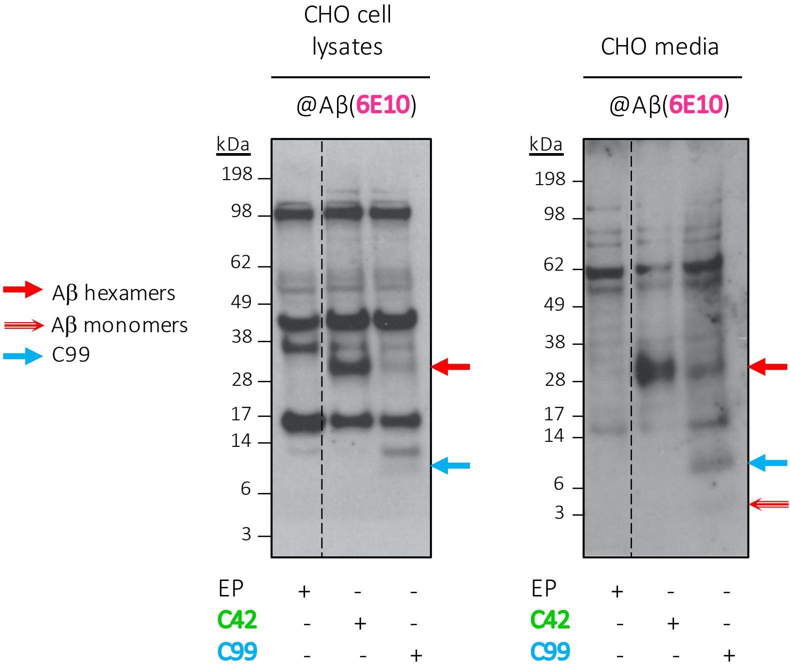
